# A Sequence Motif Enables Widespread Use of Non-Canonical Redox Cofactors in Natural Enzymes

**DOI:** 10.1101/2025.08.01.668186

**Authors:** Samer Saleh, Ning-Hsiang Hsu, Emma Luu, Vincent C. Martin, Hans Jefferson C. Ng, William B. Black, Jin-Kwang Kim, Banumathi Sankaran, Ryan L. Hayes, Justin B. Siegel, Feng Qiao, Han Li

## Abstract

Non-canonical redox cofactors (NRCs) are promising alternatives to nicotinamide adenine dinucleotide (phosphate) (NAD(P)^+^) for biomanufacturing due to low cost and exquisite electron delivery control, yet their adoption is limited by the scarcity of compatible enzymes. Here, we screened the aldehyde dehydrogenase (ALDH) protein family and identified a conserved RH/QxxR sequence motif that enables widespread NRC activity among natural enzymes. *Bos taurus* ALDH3a1 and *Pseudanabaena biceps* ALDH exhibit unprecedented turnover with nicotinamide mononucleotide (NMN^+^), with k_cat_ values matching or exceeding that of NAD^+^ and surpassing most engineered NRC-active enzymes by 10 to 10^5^-fold, based on the relative NRC to native activity. Structural and dynamic analyses reveal this motif reinforces cofactor positioning and pre-organizes the active site without dependence on the adenosine monophosphate moiety of NAD^+^. When introduced into diverse ALDH scaffolds, the RH/QxxR motif enhances NMN^+^ activity up to 60-fold. In addition to NMN^+^, this motif also supports activity across multiple non-nucleotide, simple synthetic NRCs such as 1-(2-carbamoylmethyl)nicotinamide (AmNA^+^). These findings elucidate Nature’s solution to the engineering challenge of obtaining NRC-active enzymes and offers a blueprint to mine latent evolutionary plasticity in natural enzymes that serve as superior engineering starting points.

## INTRODUCTION

Biomanufacturing, a paradigm where cell and enzyme catalysts facilitate the production of fuels, commodities, materials, medicine, and foods, is a key component in human society’s sustainable future. While biomanufacturing has been successfully applied towards a variety of products, many of these applications depend on the natural redox cofactors nicotinamide adenine dinucleotide (phosphate) (NAD(P)^+^) to mediate oxidation and reduction reactions. Inside whole cell systems, dependence on natural redox cofactors poses many challenges, including undesirable side reactions, disruption of cell fitness, and insufficient driving force for thermodynamically challenging reactions^1–3^. In cell-free systems, NAD(P)^+^ also introduces a prohibitive cost in large scale, even with recycling methods^4,5^.

Non-canonical redox cofactors (NRCs) are biomimetic analogs of NAD(P)^+^ which can replace NAD(P)^+^ in oxidoreductases^6–13^. These cofactors have garnered interest as catalysts for biotechnological purposes because they can overcome common drawbacks observed when utilizing natural cofactors as mentioned above. Nonetheless, NRCs have not yet seen wide scale adoption due to the scarcity of enzymes that can accept them effectively^13^. Engineering NRC-dependent enzymes has seen varying degrees of success, usually yielding variants that are many orders of magnitude lower in catalytic efficiency for the desired NRC compared to the natural enzyme’s native NAD(P)^+^ activity (see below).

This gap in performance between natural and engineered enzymes is well documented, which underscores the power of natural evolution in shaping enzymes^14^. In particular, designing the complex conformational dynamics required for optimal catalysis in enzymes has been recognized as a long-standing challenge, as evident by the severely suboptimal catalytic rates k_cat_ (compared to substrate binding affinity associated *K*_M_) in engineered enzymes^14^. In this study, we took a systems approach to survey a broad natural sequence space to identify NRC utilizing enzymes, distilled a sequence motif that dictates this activity, and illustrated its molecular mechanism. We envision that these enzymes not only offer ready-to-optimize scaffolds for further engineering but also teach guiding principles that can accelerate the development of efficient NRC dependent enzymes.

We chose nicotinamide mononucleotide (NMN^+^) as the model NRC for this study due to its established use in biomanufacturing, both *in vivo* and *in vitro*^8,15–19^. NMN^+^ is a biomimetic cofactor similar to NAD^+^, lacking the adenosine monophosphate (AMP) moiety (Fig. 1). Aside from flavin dependent enzymes, which are well-known to accept divergent electron donors and acceptors^20–23^, examples of NMN^+^ utilizing natural enzymes are very rare^24,25^. We selected the aldehyde dehydrogenase (ALDH) protein family (pFAM) because this large class of enzymes is diverse in sequence and function, while maintaining a high degree of structural homology^26^. This makes it easier to deconvolute sequence-function relationships and allows NRC-enabling features to be readily translated across scaffolds, as indeed demonstrated here. ALDHs have also seen use in the production of bioplastics, flavor and fragrance compounds, and biofuels, and therefore represent an interesting target for enhanced industrial biocatalysis^27–31^.

**Figure 1:**
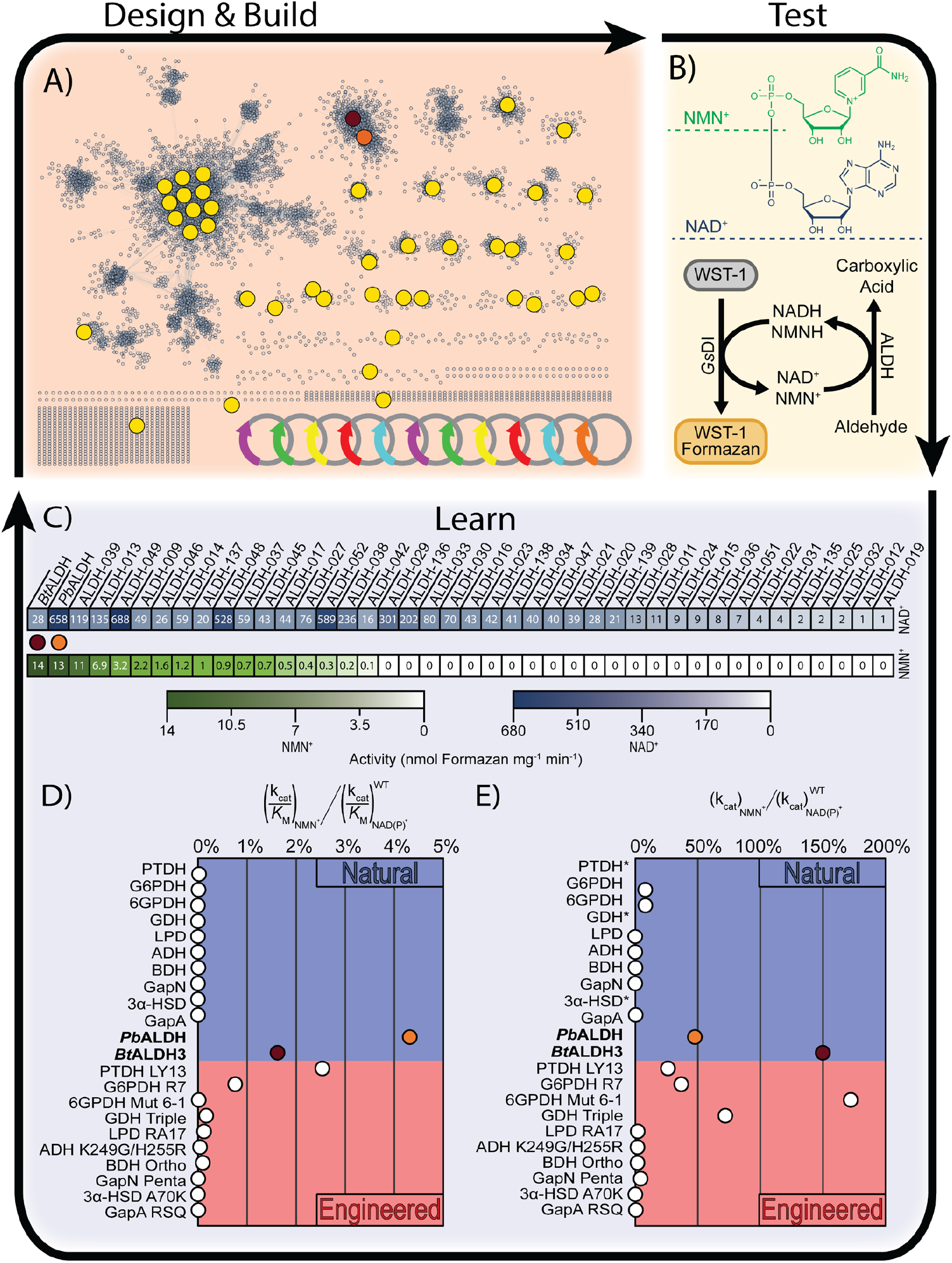
Design, build, test, and learn cycle for ALDH bioprospecting towards NMN^+^. A) The ALDH protein family sequence similarity network is used to identify and efficiently screen ALDH scaffolds. Each node represents a UniRef50 cluster of related ALDH’s, and edges represent shared sequence similarity between each node. 42 nodes (yellow, burgundy, and orange) were chosen for characterization, cloned and expressed. B) Once designed and built, each ALDH scaffold was tested in a high-throughput colorimetric cycling assay towards NAD^+^ and NMN^+^. Cofactor recycling, and WST-1 dye enable screening of many conditions and generates a sensitive readout for NMN^+^ activity with a wide dynamic range^12^. C) Results of colorimetric cycling assay; Each ALDH was assayed with its preferred substrate (acetaldehyde, butyraldehyde, or hexanal) for activity with NAD^+^ (blue) and NMN^+^ (green). The highest NMN^+^ utilizing ALDHs, *Bos taurus* ALDH3a1 (*Bt*ALDH3, burgundy node) and a putative ALDH from *Pseudanabaena biceps* (*Pb*ALDH, orange node) show remarkably high levels of NRC activity for naturally occurring enzymes. D) Comparing their ratio of catalytic efficiency (k_cat_/*K*_M_) with NMN^+^ to that with NAD^+^, these ALDHs outperform all other naturally occurring enzymes similarly characterized in the literature, and all but one engineered enzyme (phosphite dehydrogenase PTDH LY13^18^), based on their ratio relative to native cofactor (NAD(P) ^+^) activity by several orders of magnitude. E) Similarly, comparing their k_cat_ with NMN^+^ to their native cofactor k_cat_, *Bt*ALDH3 and *Pb*ALDH outperform all other naturally occurring enzymes, and most engineered enzymes in the literature. All colorimetric assay measurements were carried out with *n* = 2 biologically independent replicates.

Briefly, we generated a sequence similarity network encompassing the entire ALDH pFAM and tested representatives for activity towards NAD^+^ and NMN^+^ with high-throughput experiments. Surprisingly, more than a third of the ALDHs tested showed activity towards NMN^+^. Using a design, build, test, and learn (DBTL) approach, a signature RH/QxxR motif was discovered to be conserved in the sequence subnetwork housing the two most NRC active enzymes; *Bos taurus* ALDH3a1 (*Bt*ALDH3; UniParc: UPI00005BF137) and a putative ALDH from *Pseudanabaena biceps* (*Pb*ALDH; UniProt: L8N0N6). *Bt*ALDH3 shows 1.5-fold higher k_cat_ towards NMN^+^ (2.1 ± 0.1 s^−1^) than NAD^+^ (1.4 ± 0.1 s^−1^). *Pb*ALDH also exhibits a high k_cat_ towards NMN^+^ (3.02 ± 0.01 s^−1^). To provide a reference point, ALDHs’ functions belong to secondary metabolism. Among diverse secondary metabolism enzymes, the median value of k_cat_ for natural reactions is ~2.5 s^−1[32]^. Furthermore, *Bt*ALDH3 and *Pb*ALDH are ~10 to 10^5^-fold better than all other natural *or* engineered enzymes reported so far, based on the ratio of NMN^+^ catalytic efficiency (k_cat_ /*K*_M_) to the wild type enzyme’s native NAD(P)^+^ activity, with a single exception (engineered phosphite dehydrogenase PTDH LY13^18^).

This unusually high level of activity calls for mechanistic study. We revealed the mechanism of the RH/QxxR motif as an active site organization center using mutational studies, crystallography, and molecular dynamics modeling. The three key residues in this motif are proposed to control the conformational flexibility of the active site and reinforce cofactor positioning during catalysis via supporting interactions to an “aromatic lid”. Importantly, NMN^+^ catalytic efficiency was successfully increased by up to 60-fold when RH/QxxR was translated on to three unrelated ALDHs. RH/QxxR motif-containing ALDHs have also been shown to accept a broad range of structurally divergent, non-nucleotide, simple synthetic NRCs such as 1-(4-carboxy)benzylnicotinamide (BANA^+^), 1-(3-(4-methoxyphenyl))propylnicotinamide (P3NA-OMe^+^), and 1-(2-carbamoylmethyl)nicotinamide (AmNA^+^), opening new avenues to use these cost-effective NAD^+^ alternatives in biotransformation.

NAD(P)^+^ serve as the universal redox cofactors across all domains of life. They are widely regarded as molecular fossils of the ancient RNA world, proposed to predate even the emergence of enzymes themselves. Consequently, one may posit that redox enzymes have been evolutionarily optimized to bind NAD(P)^+^ with exquisite specificity. This work unveils that there is still considerable residual plasticity in cofactor recognition landscape, likely within enzymes that more firmly contact the electron transfer “head” of the cofactor, nicotinamide, while relying less heavily on interactions to the rest of the cofactor, which would be the “tail” that engineers seek to diversify^33^. A similar approach can be taken for other enzyme classes to understand the generalizability of this observation. Natural evolution, with its immense combinatorial depth, has generated a rich library of enzyme scaffolds that could be mined to advance NRC enzyme engineering.

## RESULTS

### High-Throughput Screening of ALDHs for Activity Towards NMN^+^

To guide our efforts in identifying NMN^+^ active ALDHs we followed a DBTL cycle (Fig. The ALDH protein family (PF00171) represents over 508,000 sequences, covering divergent functions such as detoxification of endogenous and exogenous aldehydes, synthesizing cell signaling messengers such as retinoic acid, and mitigating ocular damage from ultraviolet irradiation^26,34^. To efficiently explore the natural sequence diversity of ALDHs with experimentally tractable throughput, we reduced the redundancy of available sequences and identified representatives from each region of sequence space. To achieve this, we first reduced the number of available sequences down to ~10,000 by searching through UniRef50 cluster representatives (representatives from clustered groups of protein sequences with ≥50% identity in the UniProtKB database; see Methods for details), rather than all available sequences^35^. Next, using the UniRef50 representative sequences, we constructed a sequence similarity network (SSN) to visualize the relationships between ALDHs in sequence space (Fig. 1A)^36^.

From the ALDH SSN we chose 42 representative sequences, sampling from all the largest subnetworks, as well as a set from the isolated nodes. Sequences were primarily chosen from each subnetwork at random, however a small subset was chosen based on literature reports of attractive properties such as thermostability, high turnover, or broad substrate scope. Subnetworks containing sequences encoding for bifunctional, CoA-acylating, or phosphorylating ALDHs were avoided. The 42 chosen ALDHs originate from a wide range of prokaryotic and eukaryotic organisms and share an average of 18% pairwise sequence identity. The chosen ALDH’s were cloned, expressed in *Escherichia coli*, and purified (Fig. 1B, Tables S1, S2). The purified enzymes were then subjected to a 96-well plate colorimetric cycling assay (Fig. 1B), which utilizes a diaphorase from *Geobacillus sp*. (*Gs*DI) to rapidly oxidize the reduced cofactor (NADH or NMNH) generated by ALDH, and concomitantly reduce the WST-1 tetrazolium dye to WST-formazan, producing a sensitive readout by measuring absorbance^12^.

Since most of the chosen ALDH sequences are only inferred to belong to the ALDH pFAM by sequence homology and are otherwise entirely uncharacterized, it was necessary to perform preliminary screening to determine their preferred aldehyde substrate when coupled to NAD^+^. We selected various aliphatic aldehyde substrates (acetaldehyde, butyraldehyde, and hexanal) and coupled each ALDH with NAD^+^ to determine their substrate preference (Table S3). Next, each ALDH was assayed with its preferred aldehyde substrate and NMN^+^ as the cofactor (Fig. 1B). Surprisingly, we identified that more than a third of the ALDH’s tested had detectable activity with NMN^+^ (Fig. 1C, Table S3). The ALDH’s showing the greatest activity towards NMN^+^ were *Bt*ALDH3 and a putative ALDH *Pb*ALDH.

### Characterization of NMN^+^-Active ALDHs

We further characterized *Bt*ALDH3 and *Pb*ALDH by measuring their apparent kinetic parameters with NAD^+^ and NMN^+^ (Table 1). To provide context for the unusual levels of NMN^+^ dependent activity these enzymes exhibit, we compared these ALDHs to both natural and engineered enzymes characterized so far for NMN^+^ utilization^8,12,15,16,18,19,37–40^. From the existing literature, all other wild type enzymes show NMN^+^ activities that are 10^3^ to 10^6^ lower than their native activities with their preferred natural cofactors (NAD(P)^+^) as measured by the ratio of catalytic efficiency (k_cat_ /*K*_M_)NMN^+^ to (k_cat_ /*K*_M_)NAD(P)^+^ (Fig. 1D, Table 1). After engineering via structure-guided design or directed evolution, many of the reported enzymes showed significantly increased catalytic efficiency for NMN^+^, albeit still 10^1^ to 10^5^-fold lower than their natural activities as calculated by the ratio between (k_cat_ /*K*_M_)NMN^+^ of engineered enzymes to (k_cat_ /*K*_M_)NAD(P)^+^ wild type enzymes (Fig. 1D, Table 1). Remarkably, *Bt*ALDH3 and *Pb*ALDH, two wild type enzymes without any engineering, already have NMN^+^ catalytic efficiency that is ~2% and ~4% that of NAD^+^, respectively (Fig. 1D). This number may seem small, however, with a single exception (engineered phosphite dehydrogenase LY13, which has NMN^+^ activity ~2.5% of wild type enzyme’s natural NAD^+^ activity^18^), *Bt*ALDH3 and *Pb*ALDH are ~10^1^ to 10^5^-fold better than all other enzymes, both pre-*and* post-engineering, in terms of their (k_cat_ /*K*_M_)NMN^+^ compared to their evolutionarily honed native activity (Fig. 1D).

**Table 1:**
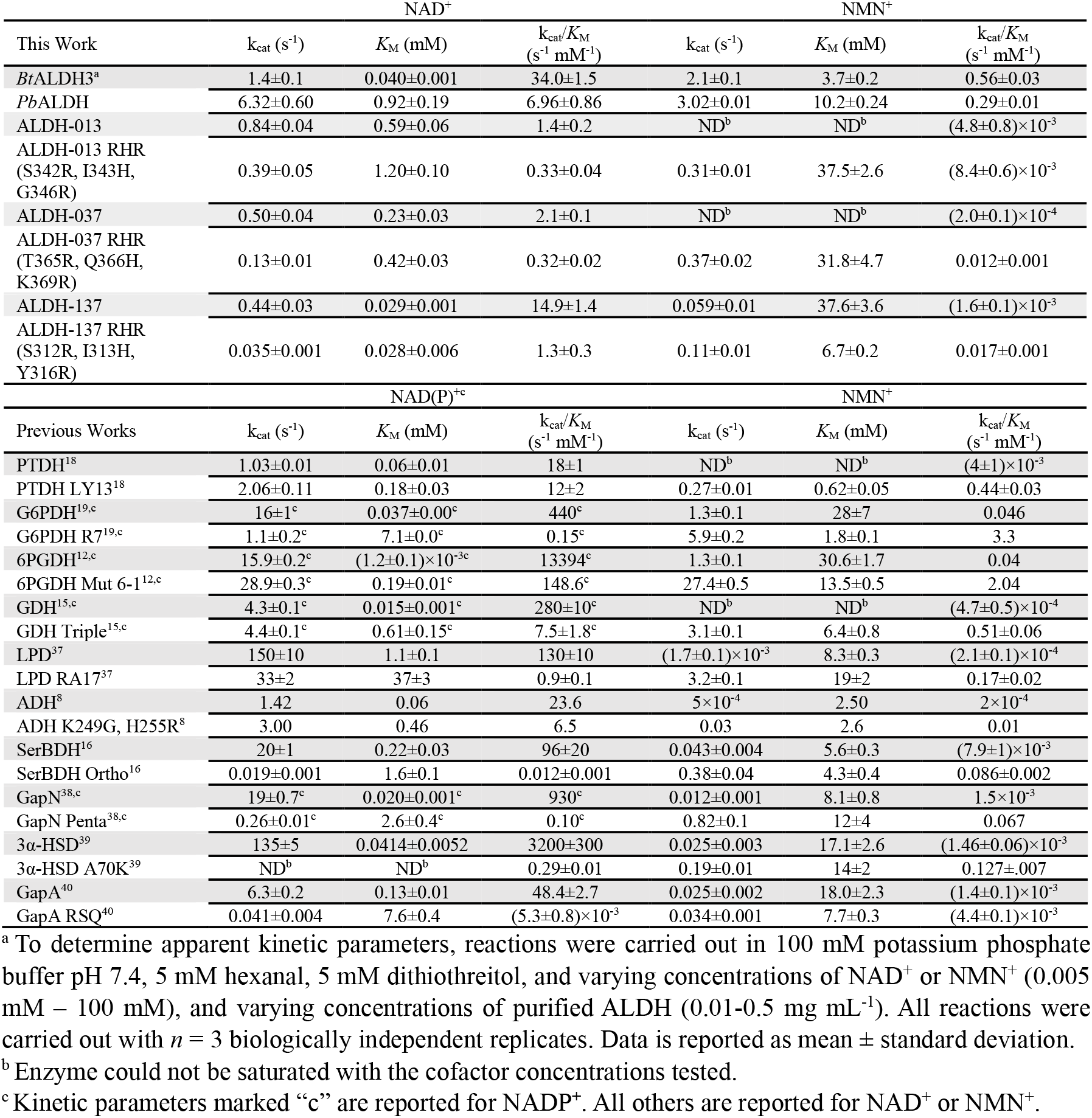
Kinetic parameters of ALDHs and NMN^+^ utilizing dehydrogenases.

While informative, using catalytic efficiency (k_cat_ /*K*_M_) as a metric might unjustly exaggerate the gap between NMN^+^ and NAD(P)^+^ activity, because *K*_M_ values strongly depend on ligand sizes^32^. Specifically, below the molecular weight cutoff of 350 Da (molecular weight of NMN^+^ = 334 Da, molecular weight of NAD^+^ = 663 Da), *K*_M_ exponentially increases as the ligand molecular weight decreases, suggesting that enzymes face intrinsic physical and chemical limitations in recognizing ligands < 350 Da^32^. These limits may be insurmountable even with natural evolution’s unparalleled depth in searching protein sequence space. Indeed, when we used k_cat_ as the metric instead, the gap between NMN^+^ and NAD(P)^+^ activity narrowed (Fig. 1E, Table 1). With a single exception (engineered 6-phosphogluconate dehydrogenase Mut6-1^12^), wild type *Bt*ALDH3 is better than all other enzymes, both pre-*and* post-engineering, in terms of the ratio between (k_cat_)NMN^+^ and the corresponding wild type enzyme’s natural (k_cat_)NAD(P)^+^ for its preferred cofactor. *Pb*ALDH also ranked high, surpassing all but two engineered enzymes in this analysis (Fig. 1E, Table 1), and outperformed wild type enzymes for which this metric can be obtained from literature.

It is important to note that the analysis above could be more robust with a broader data set on natural enzyme’s NMN^+^ kinetic parameters. Nevertheless, based on all available literature at time of publishing, our results here support that we have found superior NMN^+^ utilizing enzymes that naturally exist, which rival extensively engineered enzymes.

### Elucidating the Sequence Determinant for NMN^+^ Activity

Both *Bt*ALDH3 and *Pb*ALDH originate from the same subnetwork in the ALDH pFAM SSN (Fig. 1A). These results indicated that this subnetwork of ALDHs may be carrying a common feature in sequence space that enables activity towards NMN^+^. To investigate this hypothesis, we iterated on our DBTL approach and generated a refined SSN by isolating the nearest neighbors of *Bt*ALDH3 and *Pb*ALDH (Fig. 2A). From this refined SSN, *Bt*ALDH3 and *Pb*ALDH were found to be related by three degrees of separation, represented by five ALDH sequences. These five ALDHs include *Homo sapiens* ALDH3a1 (*Hs*ALDH3) as well as four putative ALDHs (ALDH-53, -54, -140, and -141) originating from a diverse group of prokaryotic and eukaryotic organisms and share an average of ~50 % pairwise sequence identity. All five ALDHs were cloned, expressed, and tested towards various aliphatic aldehydes (acetaldehyde, butyraldehyde, and hexanal) using NAD^+^ in a 340 nm spectrophotometric assay, and their preferred substrate was identified. Next, the specific activity of *Bt*ALDH3, *Pb*ALDH, and their five related ALDHs were similarly measured for both NAD^+^ and NMN^+^ (Fig. 2B, S1). Importantly, all five related enzymes to *Bt*ALDH3 and *Pb*ALDH demonstrated activity towards NMN^+^, and all but one (ALDH-141) showed high levels of activity towards NMN^+^. These results validate that the region of sequence space containing *Bt*ALDH3 and *Pb*ALDH in the SSN represents a group of NMN^+^ utilizing enzymes. Multiple sequence alignment of these ALDHs (Fig. 2C) uncovered a R289, Q/H290, R293 (*Bt*ALDH3 numbering) motif (RH/QxxR motif) which is conserved in the *Bt*ALDH3 sequence subnetwork but not commonly observed outside this subnetwork of ALDH sequences (as discussed below).

**Figure 2:**
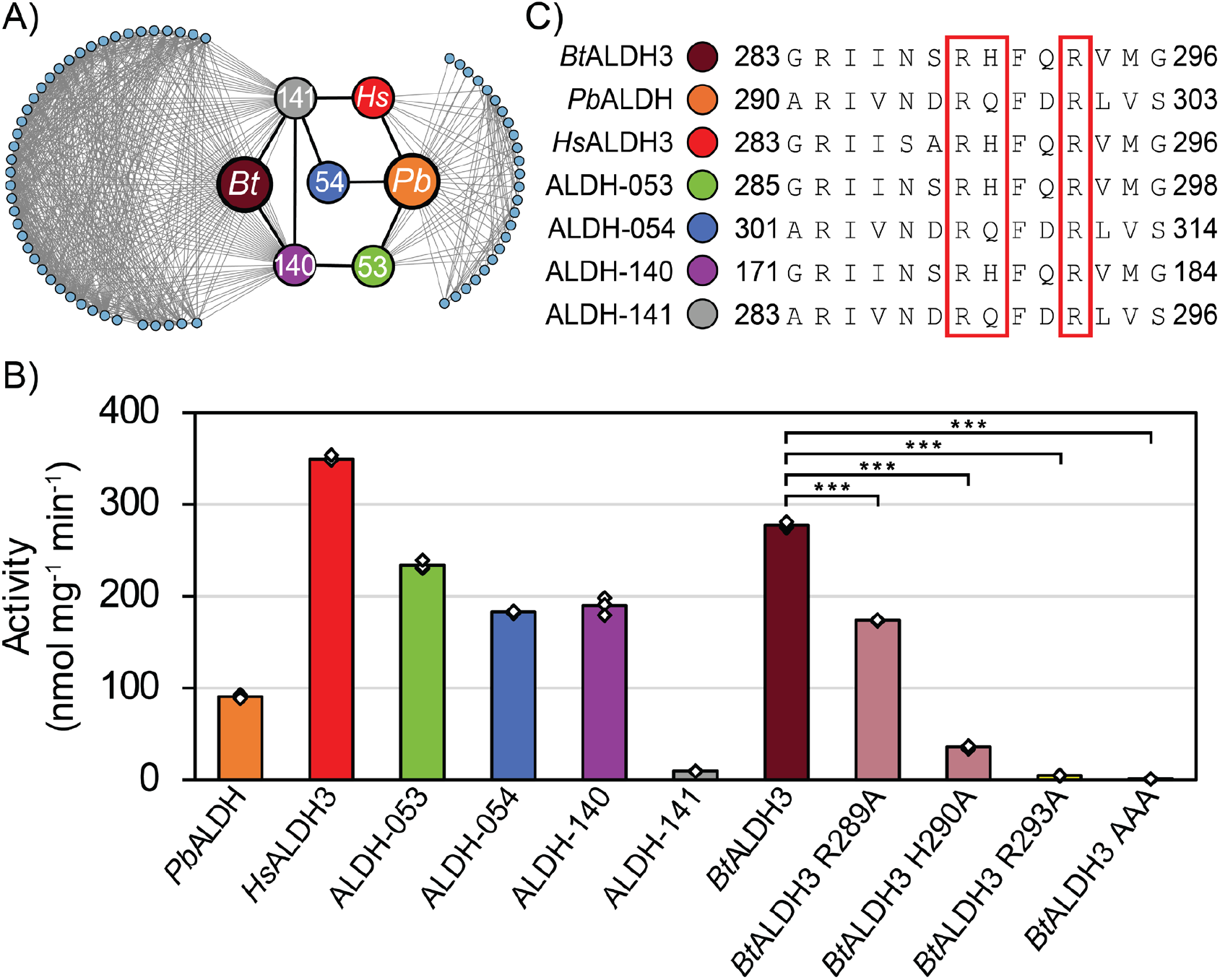
Refinement and investigation of the *Bt*ALDH3/*Pb*ALDH subnetwork. A) *Bos taurus* ALDH3a1 (*Bt*ALDH3, burgundy) and *Pseudanabaena biceps* ALDH (*Pb*ALDH, orange) show the highest level of activity towards NMN^+^ and appear in a shared region of the ALDH sequence similarity network. To investigate the sequence-function relationship of these enzymes’ NRC activity, this region was isolated to generate a refined subnetwork of their interrelated sequences. *Bt*ALDH3 and *Pb*ALDH are related by three degrees of separation encompassed by five ALDH’s sharing an average pairwise sequence identity of ~50%. B) All ALDH’s interrelated between *Bt*ALDH3 and *Pb*ALDH showed high levels of activity towards NMN^+^ when assayed with their preferred substrate (acetaldehyde for ALDH-140, hexanal for the rest). C) Multiple sequence alignment of *Bt*ALDH3, *Pb*ALDH, and their related ALDH’s uncover a highly conserved “RH/QxxR” motif unique to this subnetwork of ALDH sequences. Alanine scanning of the RH/QxxR motif on *Bt*ALDH3 reveals this motif plays a functional role in activity towards NMN^+^. “*Bt*ALDH3 AAA” corresponds to the BtALDH3 R289A, H290A, R293A triple mutant. All specific activity data were generated by monitoring the formation of reduced cofactor (NMNH) at 340 nm and was carried out with *n* = 3 independent biological replicates. Error bars correspond to one standard deviation. Statistical comparisons of WT *Bt*ALDH3 activity to *Bt*ALDH3 variants were performed using a two-tailed Welch’s t-test (*\*\*\**:*p*<0.001).

That the RH/QxxR motif is a sequence determinant of NMN^+^ activity was further supported by alanine scanning of this motif (Fig. 2B). For *Bt*ALDH3, the R289A, H290A, and R293A, single mutants showed a 47%, 89%, and 98% reduction in activity towards NMN^+^, respectively, relative to the wild type. Similarly, the *Bt*ALDH3 R289A, H290A, R293A triple mutant showed 99% reduction in activity towards NMN^+^. This established that all three key residues are important, with H290A and R293A playing the most important roles.

### Structural and Mechanistic Studies of the RQ/HxxR Motif

We solved the crystal structures of *Bt*ALDH3 with NAD^+^ and NMN^+^ bound at 1.46 Å resolution (Fig. 3A-E). Data collection and refinement statistics are listed in Table S4. *Bt*ALDH3 shares the structural architecture of class 3 dimeric ALDHs from *Rattus norvegicus*^41^, *Homo sapiens*^42^, *and Pseudomonas putida*^43^,. Each *Bt*ALDH3 structure contains two polypeptide chains per asymmetric unit which describe a homodimer. The binary complexes with NAD^+^ and NMN^+^ are defined by clear electron densities, although a minor NAD^+^ electron density is seen in the NMN^+^ complex. This is likely due to contaminating NAD^+^ carried over from purification which remains bound to a minor fraction of protein during co-crystallization and is most prominent in subunit A. The primary interactions to the cofactors are formed at the carboxamide, NMN^+^ ribose, and NMN^+^ phosphate, and are mediated by E210, E344, S189, and W114 (Fig. 3A-E). In addition to these interactions, the NAD^+^ complex shows additional interactions to the adenosine monophosphate (AMP) ribose through K138 and E141.

**Figure 3:**
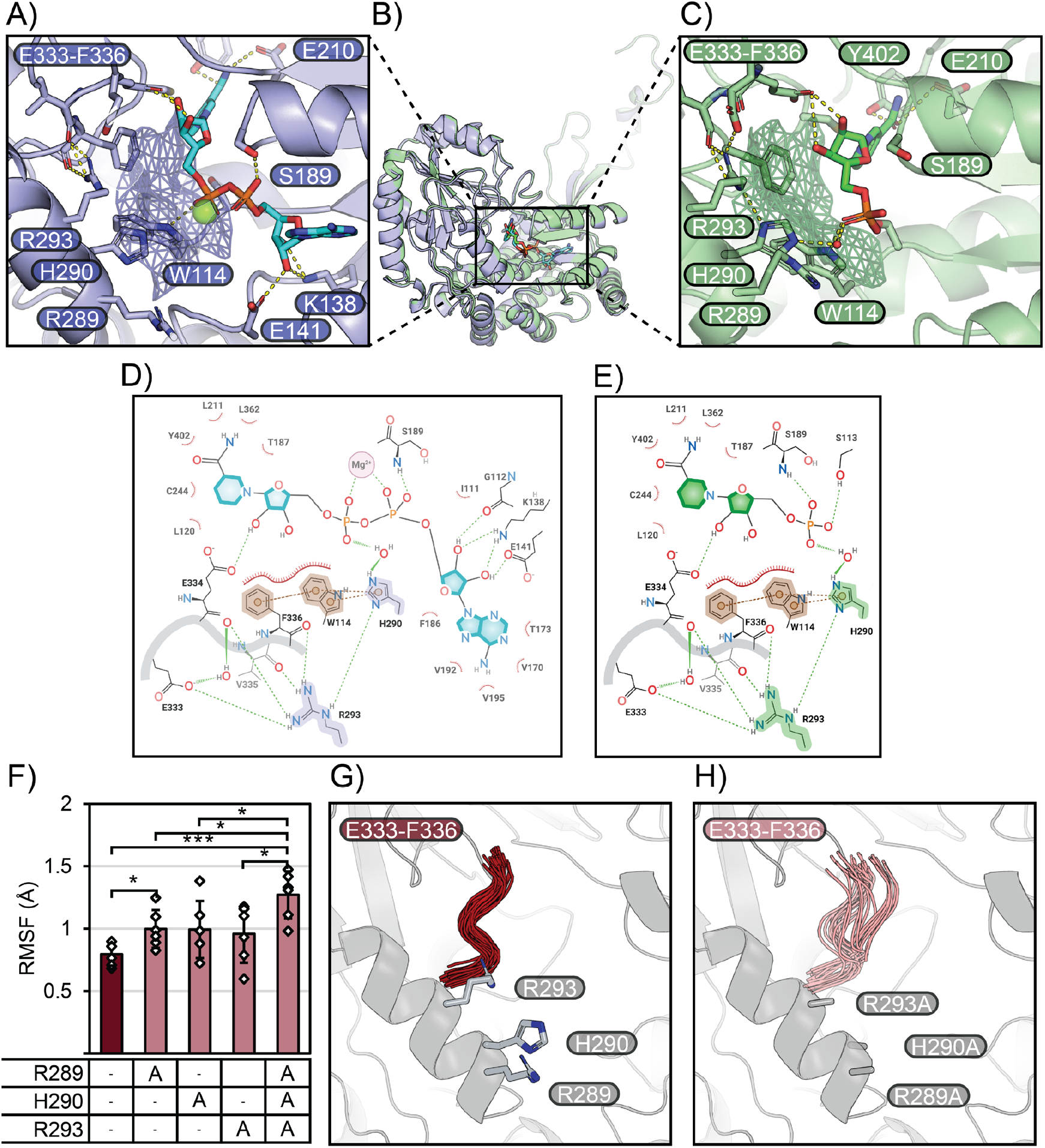
Structural and mechanistic analysis of *Bt*ALDH3. A-E) Crystal structure and interaction maps of *Bos taurus* ALDH3a1 (*Bt*ALDH3). The overall structure of *Bt*ALDH3 (B), follows the architecture of dimeric class 3 ALDHs. The cofactor binding pocket with NAD^+^ (A, D) and NMN^+^ (C, E) share the same hydrogen bond interactions through W114, S189, E210, E334, and Y402F, indicating that active site organization is independent of adenosine monophosphate (AMP) binding. The cofactors are also stabilized by an “aromatic lid” formed by W114, and F336 that pack against NMN^+^. The RH/QxxR motif of *Bt*ALDH3 is captured in the R289 (whose electron density could not be resolved), H290, R293 residues. R293 coordinates a complex hydrogen bond and salt bridge network to the E333-F336 loop, while H290 stabilizes W114, R293, and F336 through π-π stacking and hydrogen bond interactions, facilitating active site pre-organization. F) Molecular dynamics simulations of apo *Bt*ALDH3 WT and alanine scanning variants shows a significant difference in the root mean square fluctuation (RMSF) of the E333-F336 loop main-chain, between the WT (0.79 ± 0.09 Å) and the R289A, H290A, R293A triple mutant (1.27 ± 0.19 Å, *p* < 0.001). Single mutants of *Bt*ALDH3 show similar increases in RMSF: R289A (1.00 ± 0.15 Å, p < 0.05), H290A (0.99 ± 0.23 Å, p = 0.09), and R293A (0.96 ± 0.23 Å, p = 0.15). For visualization, thirty random frames are chosen from the WT (G) and R289A, H290A, R293A triple mutant (H) simulations. WT *Bt*ALDH3 maintains an organized loop structure through interactions to the RH/QxxR motif, while total elimination of RH/QxxR results in a disordered active site. Simulations were carried out with *n =* 6 monomer simulations. Error bars correspond to one standard deviation. Pairwise statistical comparisons between RMSF values of ALDH variants were performed using a two-tailed Welch’s t-test (*:*p*<0.05, *\*\*\**:*p*<0.001).

To our surprise, NAD^+^ and NMN^+^ bind nearly identically to *Bt*ALDH3 with a root mean square deviation of 0.14 Å for NMN^+^ portion of both cofactors. Similar to the NAD^+^ conformation captured in the crystal structures of the class 3 ALDHs from *Homo sapiens*, and *Pseudomonas putida, Bt*ALDH3 co-crystallizes with NAD^+^ and NMN^+^ in their hydride-transfer poses^42–46^. The structural architecture of *Bt*ALDH3 shows no significant differences in main-chain conformation or side-chain orientation upon NAD^+^ or NMN^+^ binding. In contrast, cofactor binding has been shown to modulate enzymes conformational dynamics in numerous enzymes^47–51^. In particular, the AMP moiety of the NAD^+^, despite being reaction inert, can exert long range allosteric effects to pre-organize the enzyme active site^24,52^. In fact, buttressing the void in enzyme’s AMP binding region has been proven as a generalizable design principle in engineering NMN(H)-dependent enzymes^16,18,53^. As discussed below, our results indicate that the RH/QxxR motif contributes to the high rigidity of *Bt*ALDH3 active site, such that catalysis is well ordered without relying on the AMP moiety of NAD^+^ to orchestrate active site organization.

Firstly, the *Bt*ALDH3 crystal structures reveals that F336 and W114 form an “aromatic lid” (Fig. 3A, C) that encloses NMN^+^ in the binding pocket, potentially supporting this non-canonical cofactor through catalysis without AMP anchoring. Notably, the RH/QxxR motif is critical for positioning this “aromatic lid”. R293 forms a series of hydrogen bonds to the backbones of E334, V335, and F336 as well as a salt bridge with E333 (Fig. 3C, E). This network of interactions nucleated by R293 reinforces the conformation of the loop spanning E333-F336, which holds F336 in place. H290 engages in π–π stacking with W114, which further stacks with F336. Additionally, H290 forms a supporting hydrogen bond that coordinates R293, and a water mediated interaction to the NMN^+^ phosphate (Fig. 3C, E).

The electron density of R289 is not well resolved, indicating this residue is solvent exposed and highly flexible. We hypothesize that it modulates the electrostatic environment and supports the hydrogen bond network mediated through H290 and R293. This mechanism, though not captured in the static crystal structure, was inferred from molecular dynamic studies (see below). The spacer residues “xx” in the RH/QxxR motif are required to maintain the α-helix between H290 and R293, positioning R293 to face the active site and interact directly with the E333-F336 loop. While their side-chain identities are relatively conserved among the natural NRC utilizing ALDHs tested above, those side-chains are solvent exposed and do not appear to have any notable synergy with the rest of the motif. Ultimately, while the R289, H290, and R293 are required for enabling NRC activity, the side-chains identities “xx” can be variable (see below).

Secondly, we performed molecular dynamic analysis to pinpoint the effect of the RH/QxxR motif in stabilizing the catalytically important E333-F336 loop. Besides housing the “aromatic lid” residue F336, the E333-F336 loop may have a broader impact on active site dynamics. F336 is responsible for modulating the conformational dynamics of the nicotinamide portion of the cofactor during catalysis^44^. Moreover, the E333-F336 loop is adjacent flexible loop containing the central catalytic residue of *Bt*ALDH3, C244. Therefore, fastening down the E333-F336 loop may clear clashes and excessive fluctuations to protect catalytic events from interference. Dynamics simulations on apo WT *Bt*ALDH3, as well as apo R289A, H290A, R293A single mutants, and the corresponding triple mutant were conducted. The flexibility of E333-F336 was assessed by the average root mean square fluctuation (RMSF) across the alpha-carbons of E333-F336 after alignment to the average trajectory structure (Fig. 3F). The triple alanine mutation significantly increases the RMSF of the E333-F336 loop from 0.79 ± 0.09 Å to 1.27 ± 0.19 Å (*p* < 0.001). This can be visually understood by observing that the E333-F336 loop adopts an ordered conformation in WT *Bt*ALDH3 (Fig. 3G) but adopts a wider range of catalytically irrelevant poses upon mutating the RH/QxxR motif (Fig. 3H). Similar increases in the RMSF were observed for the single mutants R289A (1.00 ± 0.15 Å, *p* < 0.05), H290A (0.99 ± 0.23 Å, *p* = 0.09), and R293A (0.96 ± 0.23 Å, *p* = 0.15), compared to the WT, but only the R289A mutant showed a statistically significant increase. These results suggest the full RH/QxxR motif is the most effective in organizing the active site; when studied in the context of dynamic structural behavior, R289 plays a significant role in the RH/QxxR motif.

### The Effect Of RH/QxxR Motif Is Transferrable to Diverse ALDHs and NRCs

To test the extent to which the RH/QxxR motif is self-sufficient in enabling NMN^+^ activity, we analyzed the ALDH pFAM SSN for subnetworks whose consensus sequences do not contain this motif (Fig. 4A, 4B). From these, we chose three phylogenetically distinct ALDHs (ALDH-013, -037, -137) to mutate at their equivalent positions and replicate the RH/QxxR motif, leaving the “xx” positions as their respective wild type identities. The RH/QxxR mutants of ALDH-013 (S342R, I343H, G346R), ALDH-037 (T365R, Q366H, K369R), and ALDH-137 (S312R, I313H, Y316R) scaffolds showed 2.5 to 18-fold improvements in specific activity towards NMN^+^ over their WT activities (Fig. 4C). Kinetic characterization showed that ALDH-013, ALDH-137, and ALDH-037 mutants showed a 2-, 60-, and 8-fold increase in catalytic efficiency towards NMN^+^, respectively, compared to their respective wildtypes (Table 1). These improvements are primarily driven by reductions in the *K*_M_, which supports that the RH/QxxR motif enhances NMN^+^ binding by rigidifying and pre-organizing the cofactor binding pocket. Accordingly, the RHR mutants did not enhance catalytic efficiency toward NAD^+^.

**Figure 4:**
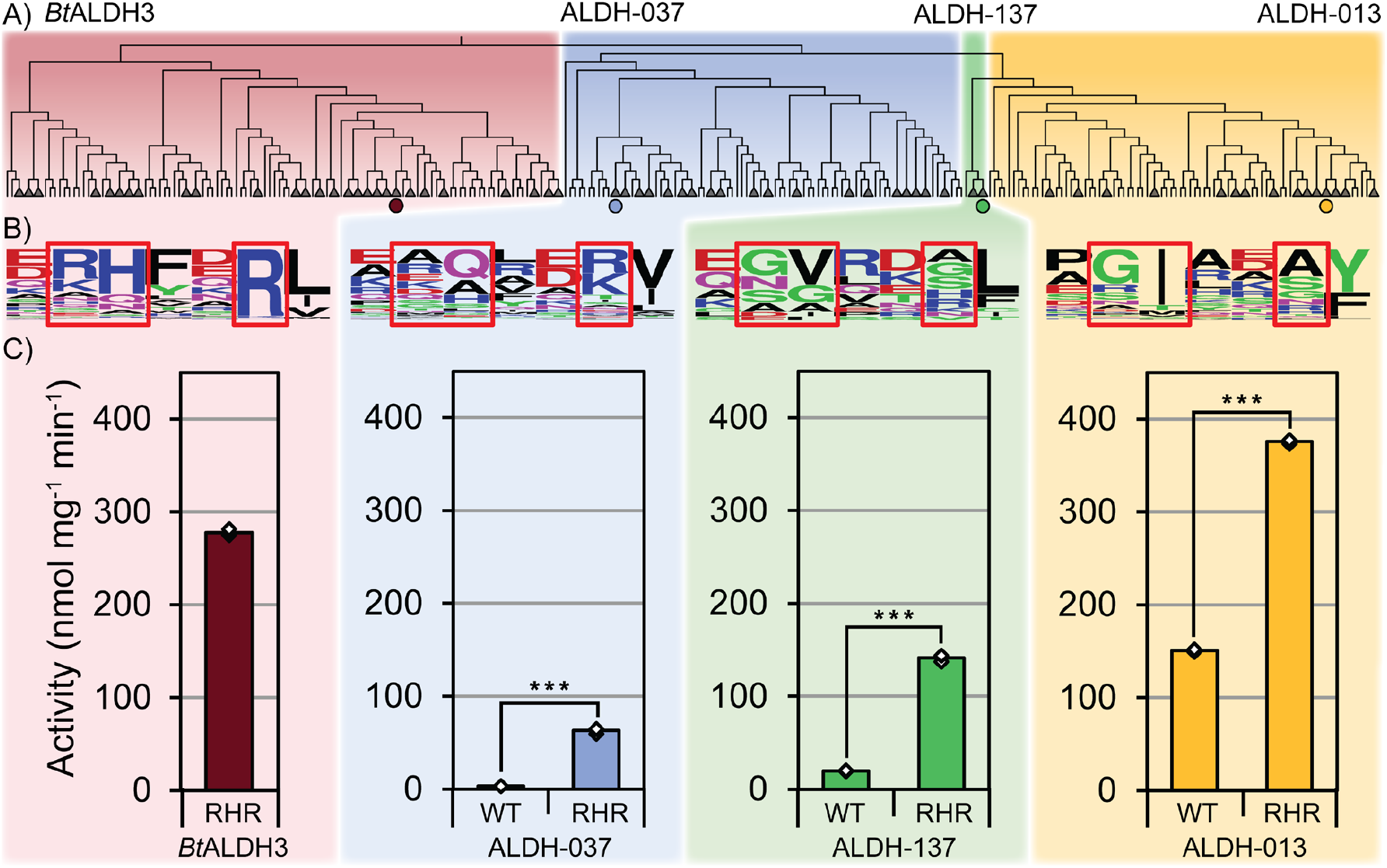
Transferring of RH/QxxR motif from *Bt*ALDH3 onto ALDH-037, -137, and -013. A) To investigate the transferability of the RH/QxxR motif, three ALDH scaffolds with varying levels of activity towards NMN^+^ were chosen from phylogenetically and sequence distinct subnetworks of the ALDH protein family sequence similarity network. B) Of all the ALDH protein family subnetworks, only the *Bos taurus* ALDH3a1 subnetwork carries the RH/QxxR motif as the consensus sequence. C) Upon transferring the RH/QxxR motif onto ALDH-037 (T365R, Q366H, K369R), ALDH-137 (S312R, I313H, Y316R), and ALDH-013 (S342R, I343H, G346R) we observed an 18-, 7-, and 2.5-fold increase in activity towards NMN^+^, respectively, demonstrating the RH/QxxR motif’s central role in enabling NRC activity. All reactions were carried out with *n* = 3 biologically independent replicates. Error bars correspond to one standard deviation. Statistical comparisons between WT and “RHR” variants were performed using a two-tailed Welch’s t-test (*\*\*\**:*p*<0.001).

While NMN^+^ has many advantages as an NRC, other non-nucleotide, simple synthetic cofactors have also garnered interest due to their facile synthesis, reduced cost, and enhanced redox potential^6,9,10,11,20–23,53^. We theorized that the RH/QxxR motif may also stabilize the binding of other NRCs, which vary in size and polarity, but all share the nicotinamide “head” with NMN^+^ that the “aromatic lid” packs against (Fig. 5A). In addition to testing *Bt*ALDH3, *Pb*ALDH, and *Hs*ALDH3 we chose to test the well characterized class 3 ALDH from *Pseudomonas putida* (*Pp*MdlD), which also contains the RH/QxxR motif (R290, Q291, R294). *Pp*MdlD is a promising industrial catalyst owing to its high thermostability, broad substrate scope, and high turnover rates^43^. First, we synthesized the six non-nucleotide, simple synthetic cofactors 1-benzylnicotinamide (BNA^+^), 1-(4-carboxy)benzylnicotinamide (BANA^+^), 1-phenethylnicotinamide (P2NA^+^), 1-(3-phenyl)propylnicotinamide (P3NA^+^), 1-(3-(4-methoxyphenyl))propylnicotinamide (P3NA-OMe^+^), and 1-(2-carbamoylmethyl)nicotinamide (AmNA^+^). Next, we performed a biotransformation assay over the course of 24 hours and measured the cofactor dependent conversion of 20 mM benzaldehyde to benzoate mediated by the four chosen ALDHs. Excitingly, all enzymes tested showed conversion using simple synthetic NRCs (Fig. 5B). *Pb*ALDH showed the highest level of conversion with all the simple synthetic cofactors tested, achieving >5 mM (>25%) conversion with all cofactors, and up to 16.3 ± 0.1 mM (81.5%) conversion with AmNA^+^. Similarly, *Pp*MdlD showed conversion with all cofactors tested, albeit at lower levels than *Pb*ALDH, achieving up to 9.9 ± 0.2 mM (49.4 %) conversion with AmNA^+^. It appears that the more polar cofactors (BANA^+^, AmNA^+^, P3NA-OMe^+^) were more readily accepted as compared to their non-polar counterparts (BNA^+^, P2NA^+^, P3NA^+^), consistent with the trend observed for natural enzyme-ligand interaction^32^.

**Figure 5:**
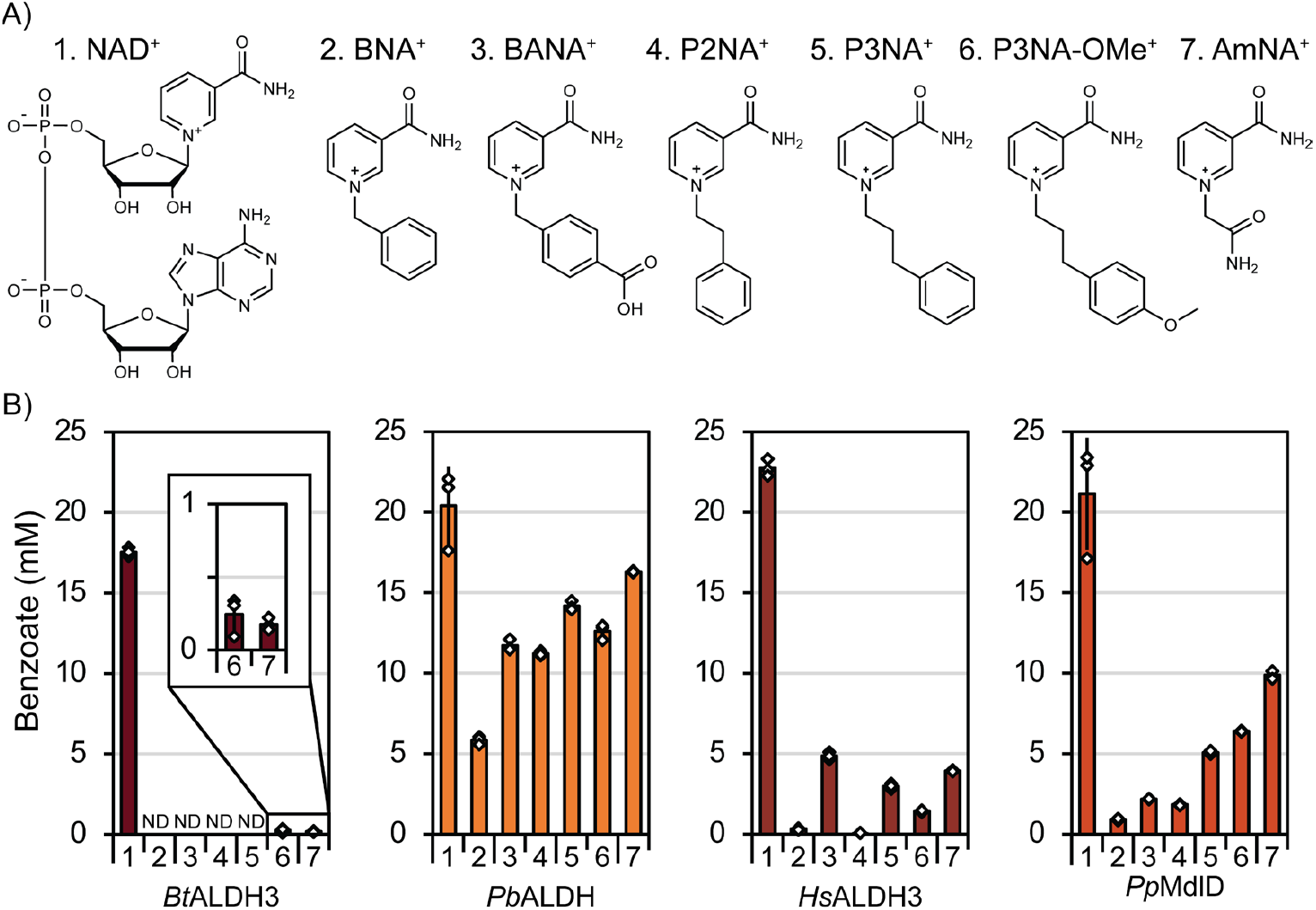
RH/QxxR motif enables NRC activity with non-nucleotide, simple synthetic NRCs. A) The simple NRCs 1-benzylnicotinamide (BNA^+^), 1-(4-carboxy)benzylnicotinamide (BANA^+^), 1-phenethylnicotinamide (P2NA^+^), 1-(3-phenyl)propylnicotinamide (P3NA^+^), 1-(3-(4-methoxyphenyl))propylnicotinamide (P3NA-OMe^+^), and 1-(2-carbamoylmethyl)nicotinamide (AmNA^+^) are readily synthesized and allow functional probing of cofactor promiscuity in ALDHs. B) In a conversion assay, *Bos taurus* ALDH3a1 (*Bt*ALDH3) only shows less than 0.25 mM conversion of benzaldehyde to benzoate with simple synthetic NRCs over 24 hours. However, *Pseudanabaena biceps* ALDH (*Pb*ALDH) shows much higher levels of benzaldehyde conversion with simple synthetic NRCs, up to 16.3 ± 0.1 mM (81.5% conversion) with AmNA^+^ and 14.2 ± 0.3 mM (71% conversion) with P3NA^+^. *Homo sapiens* ALDH3 (*Hs*ALDH3) and *Pseudomonas putida* (*Pp*MdlD) which also carry the RH/QxxR motif, showed intermediate levels of conversion, generally preferring the more polar cofactors BANA^+^, P3NA-OMe^+^, and AmNA^+^. All reactions were carried out with *n* = 3 biologically independent replicates. “ND” represents no benzoate detected.

*Bt*ALDH3 only showed detectable conversion with P3NA-OMe^+^ and AmNA^+^. On the other hand, *Hs*ALDH3, which shares 83% sequence identity to *Bt*ALDH3, displays substantially higher conversion with a broader scope of cofactors, reaching up to 4.9 ± 0.2 mM (24.3 %) conversion with BANA^+^ (Fig. 5B). This is likely to be understood in the context of the relatively inferior catalytic rate constant of *Bt*ALDH3 (k_cat, NAD+_ = 1.4 ± 0.1 s^−1^) compared to *Hs*ALDH3 (k_cat, NAD+_ = 14.1 ± 0.4 s^−1^) (Table S5). This trend persists for other high conversion ALDHs *Pb*ALDH and *Pp*MdlD, both having faster k_cat, NAD+_than *Bt*ALDH3 (Table 1, S5). Enzymes have an intrinsic rhythm at which their structures fluctuate^54–56^. These vibrations which occur, even for apo enzymes, are believed to promote catalysis upon substrate encounter. Overall, our results suggest that the RH/QxxR motif can serve as a sequence indicator for searching NRC-dependent activity in ALDH enzymes beyond NMN^+^. However, the magnitude of this activity, in particular the conversion rate in biotransformation under relatively high cofactor concentrations, may be associated with the innate conformational dynamics of enzymes.

During the preparation of this manuscript a parallel effort by Zhou et al. demonstrated that an ALDH from *Sphingobium sp*. can also utilize simple synthetic cofactors (Fig. S2)^57^. *Sp*ALDH2 also contains the RH/QxxR motif (R343, Q344, R347), which represents another independent piece of evidence corroborating our findings here.

## DISCUSSION

In this work we used a high-throughput DBTL approach, iteratively leveraging the natural sequence space, to discover and engineer ALDHs towards NMN^+^, with an extension to other simpler synthetic NRCs. While previous studies have focused on engineering activity towards NRCs from the ground up based on a particular scaffold^7,8,10–13,15–19,22,23,37–40^, we systematically screened through the ALDH pFAM and identified naturally occurring NMN^+^ utilizing ALDHs. Among these, *Bt*ALDH3 and *Pb*ALDH, are ~10^1^-10^5^-fold better than all other natural or engineered enzymes reported so far, as measured by the ratio between NMN^+^ catalytic efficiency and wild type enzymes native NAD(P)^+^ catalytic efficiency. A single exception to this is phosphite dehydrogenase PTDH LY13, which reaches similar level of relative catalytic proficiency to *Bt*ALDH3 and *Pb*ALDH for NMN^+^ (Fig. 1D). Interestingly, PTDH LY13 was obtained through a cell-fitness based, *in vivo* directed evolution campaign^18^. This selection process parallels our search through natural evolutionary sequence space, albeit much smaller in scale and sequence space depth.

Using a systems approach, for the first time we identified a distinct sequence motif that determines NRC utilization in natural enzymes. Although *Bt*ALDH3 and *Pb*ALDH only share 46% sequence identity and are phylogenetically distant, we were able to use sequence similarity networks to isolate their relationship in sequence space and distill the RH/QxxR sequence motif. This motif is not only validated to be essential in *Bt*ALDH3 for NMN^+^ activity but can also be translated to other unrelated ALDHs to impart NMN^+^ activity. Biochemical, structural, and computational studies elucidated that RH/QxxR motif functions through rigidifying a loop in the active site to more firmly support the hydride transfer portion of cofactors, without discriminating different “recognition handles” of NRCs. In line with these findings we also successfully mined non-nucleotide, simple synthetic NRC activities in natural ALDHs guided by the presence of RH/QxxR.

The RH/QxxR motif’s mechanism can be classified as active site pre-organization, which has been shown to be more optimized in natural enzymes versus engineered enzymes (e.g. natural enzymes have active site residues pointing in catalytically competent directions more frequently than engineered enzymes) ^14,58^. Active site pre-organization requires more complex and integrated interaction networks as exemplified here by *Bt*ALDH3, which is challenging to rationally design, though the latest ensemble modeling and deep-learning methods may contribute new solutions to this challenge^58,59^. Directed evolution, which emulates natural evolution in microcosm, has been shown to improve enzyme function by narrowing enzyme’s conformational ensemble to enrich for catalytically competent states^60^. Through studying the mechanisms of RH/QxxR motif, it appears that NRC-utilizing ALDHs feature a narrowly defined active site shaped by evolution.

The prevalence of significant NMN^+^ activity in natural enzymes raises a fundamental question: is this activity a byproduct of non-physiological promiscuity or does it indicate some physiological relevance? One can find partial support for either hypothesis: The RH/QxxR motif appears to be a highly conserved feature of dimeric ALDHs^34,42,43^. It is possible that the motif evolved to compensate for stabilizing interactions normally mediated by additional monomers in higher-order oligomeric assemblies. As a side product, the stabilized active side gained promiscuous activity toward NRCs, which lack AMP anchor in NAD^+^ and would otherwise be dislodged by the excessively flexible loops^53^.

On the other hand, although adenosine is suggested to be the product of an RNA world that was fossilized in contemporary cofactors, Nature has exploited both adenylated and non-adenylated versions of cofactors such as flavin adenine dinucleotide (FAD) and flavin mononucleotide (FMN) for flavins or adenosine triphosphate (ATP) and pyrophosphate in energy metabolism^61,62^. Since the AMP moiety is added at the cost of ATP, evidence suggests that the simpler, non-adenylated cofactor may function under stress or energy-constrained conditions^63,64^. The RH/QxxR motif is enriched in class 3 ALDHs in higher organisms. Members of class 3 ALDHs such as *Bt*ALDH3 and *Hs*ALDH3 are primarily located in the eyes, an organ that is particularly vulnerable to oxidative stress due to constant radiation exposure^34,65^. Indeed, high oxidative stress can impair NAD^+^ biosynthesis in ocular tissues through aging and disease, leading to the accumulation of NAD^+^ precursors such as NMN^+66^. While NMN^+^ is present at significant levels in humans, and other organisms, as an NAD^+^ biosynthesis precursor, more studies are certainly required to draw any conclusion on whether this non-adenylated version of NAD^+^ is actually utilized as a redox cofactor *in vivo* by naturally occurring enzymes such as those identified in this work^67,68,69^.

## METHODS

### Generation of ALDH Sequence Similarity Network

Sequence similarity networks were generated using the EFI-EST webtool supplied with the ALDH protein family (PF00171) as the input query, filtered by UniRef50 sequences^36^. UniRef50 clusters are clustered groups of unique sequences from the UniProtKB database sharing 50% sequence identity, and 80% sequence length overlap, which are assigned a representative sequence by choosing either the highest quality UniProt entry in the cluster, sequences deriving from model organisms, or by assigning the longest sequence as the representative^35^. Generated networks were visualized with Cytoscape. The refined *Bt*ALDH3 subnetwork (Fig. 2A) was generated in Cytoscape by selecting *Bt*ALDH3 and *Pb*ALDH, extracting their nearest neighbors using the built-in selection tool, and exporting the connected subset as a new network.

### Plasmid Construction

All plasmids used in this study are listed in Table S1, with corresponding amino acid sequences in Table S2. To clone the *Geobacillus sp*. diaphorase (*Gs*DI) and ALDHs used in this study, their DNA sequences were first codon optimized for expression in *E. coli* and their corresponding gene fragments were obtained as synthesized oligonucleotides (Integrated DNA Technologies). Synthesized gene fragments were inserted into the pQElac vector (PLlacO1, ColE1 ori, AmpR, N-terminal 6x His-tag) using Gibson isothermal DNA assembly method. To isolate plasmids for sequence verification, XL1-Blue competent cells (Agilent Technologies) were transformed with the Gibson assembly products and plated on 2×YT agar plates (16 g L^−1^ tryptone, 10 g L^−1^ yeast extract, 5 g L^−1^ NaCl) containing 100 mg L^−1^ ampicillin and incubated at 37 °C overnight. Single colonies were chosen from each construct, inoculated into 5 mL of 2×YT liquid media with 100 mg L^−1^ ampicillin and shaken overnight at 37 °C. Plasmid DNA was then extracted from each culture using the QIAprep Spin Miniprep Kit (Qiagen), and sequence verified via Sanger sequencing. pET28a-His-Smt3-*Bt*ALDH3 was similarly constructed using pET28a-His-Smt3 (T7:: BtALDH3, pBR322 ori, KanR, N-terminal 10xHis-Smt3 tag) vector in place of pQElac.

Variants of *Bt*ALDH3, ALDH-013, ALDH-037, and ALDH-137 were generated through site directed mutagenesis by PCR using the KOD1 PCR Master Mix (Toyobo). Mutated gene fragments were amplified using primers containing point mutations (Integrated DNA Technologies). Gene fragments were assembled into pQElac, isolated, and sequence verified as described above.

### Protein Expression and Purification

The ALDH’s characterized in this study were expressed in *E. coli* BL21 (Invitrogen) transformed with the respective pQElac ALDH plasmid. To begin, single colonies of *E. coli* BL21, each carrying an ALDH expression plasmid, were inoculated into individual 10 mL culture tubes containing 5 mL 2×YT liquid medium containing 100 mg L^−1^ ampicillin, and shaken overnight at 30 °C. Next, 50 mL of 2×YT medium supplemented with 200 mg L^−1^ ampicillin was inoculated with 0.5 mL of overnight culture in a 250 mL baffled flask. Cultures were grown at 37 °C, 250 RPM until reaching OD ~0.6-1, then induced with 0.5 mM isopropyl β-D-1-thiogalactopyranoside (IPTG) and incubated at room temperature. After 24 hours of expression, the cultures were transferred to 50 mL falcon tubes and centrifuged at 4,000×g, 4 °C. The supernatant was discarded, and the cell pellet was resuspended in 1 mL of ice-cold equilibration buffer (50 mM sodium phosphate buffer pH 7.7, 300 mM sodium chloride 10 mM imidazole, 0.03% Triton X-100) and transferred to a pre-chilled 2 mL bead beating tube containing 0.5 mL of 0.1 mm diameter soda lime glass beads (BioSpec Products) and stored on ice. Each resuspended cell pellet was then homogenized on a benchtop homogenizer (FastPrep-24, MP Biomedicals) at 4 m s^−1^ for 35 seconds and cooled on ice for 5 minutes. This process was repeated twice for three total homogenization steps. The crude lysate was separated from the cell debris by centrifugation at 21,000×g, 4 °C and purified using HisPur Ni-NTA Protein Miniprep (Thermo Fisher Scientific) according to the manufacturer’s instructions. All enzymes were eluted in protein elution buffer (50 mM sodium phosphate buffer pH 7.7, 300 mM sodium chloride 250 mM imidazole) and stored in 20 % (v/v) glycerol at –80 °C. *Gs*DI was similarly expressed and purified, with the only deviation being post-induction incubation at 30 °C rather than room temperature.

### High-throughput Colorimetric Cycling Assay

The high-throughput colorimetric cycling assay (Fig. 1C, Table S3) was carried out as follows: A 100 μL reaction was prepared containing 100 mM potassium phosphate buffer pH 7.4, 5 mM aldehyde (acetaldehyde, butyraldehyde, or hexanal), 0.5 mM WST-1 tetrazolium dye, 3 mM cofactor (NAD^+^, NMN^+^, or “no cofactor”), and 1 μM of purified *Gs*DI, and varying concentrations of purified ALDH (0.01-0.5 mg mL^−1^). Reactions were initiated by the addition of ALDH to reaction buffer and carried out in 96-well plates at 30 °C. The formation of WST-1 formazan was monitored kinetically at 438 nm (ε_WST-1 Formazan_ = 37×10^3^ M^−1^ cm^−1^) and activity is reported as nmol WST-1 formazan generated per mg of ALDH per minute. All activity data was corrected by subtracting “no-cofactor” controls, where cofactor was substituted with DI water in the reaction mixture and the background activity was measured. All reactions were carried out with *n* = 2 biologically independent replicates.

### Specific Activity Assays

The specific activity assays (Fig. 2B, 4C) were carried out as follows: A 200 μL reaction was prepared containing 100 mM potassium phosphate buffer pH 7.4, 5 mM aldehyde (acetaldehyde, butyraldehyde, or hexanal), 5 mM dithiothreitol, and 3 mM cofactor (NAD^+^, NMN^+^, or “no cofactor”), and varying concentrations of purified ALDH (0.01-0.5 mg mL^−1^). Reactions were initiated by the addition of ALDH to reaction buffer and carried out in 96-well plates at 30 °C. The formation of reduced cofactor was monitored kinetically at 340 nm (ε_NADH/NMNH_ = 6.22×10^3^ M^−1^ cm^−1^). All activity data was corrected by subtracting “no-cofactor” controls, where cofactor was substituted with DI water in the reaction mixture and the background activity was measured. All reactions were carried out with *n* = 3 biologically independent replicates.

### Steady-State Kinetic Characterization

The steady-state kinetic characterization of ALDH’s was performed as follows: A 200 μL reaction was prepared containing 100 mM potassium phosphate buffer pH 7.4, 5 mM hexanal, 5 mM dithiothreitol, and varying concentrations of NAD^+^ or NMN^+^ (0.005 mM – 100 mM), and varying concentrations of purified ALDH (0.01-0.5 mg mL^−1^). Reactions were initiated by the addition of ALDH to reaction buffer and carried out in 96-well plates at 30 °C. The formation of reduced cofactor was monitored kinetically at 340 nm (ε_NADH/NMNH_ = 6.22×10^3^ M^−1^ cm^−1^). The resulting specific activity data was plotted and fit to the standard Michaelis-Menten equation. In cases where the enzyme could not be saturated with 100 mM cofactor, it was assumed that K_m_>>> 100 mM, and therefore the kinetic parameters were determined using the simplified Michaelis-Menten equation (V_o_≈ k_cat_ /K_m_). All reactions were carried out with *n* = 3 biologically independent replicates.

### Crystallography of *Bt*ALDH3

The plasmid pET28a-His-smt3-*Bt*ALDH3 was transformed into *E. coli* Rosetta BL21(DE3) cells. Protein expression was induced in log-phase cultures with 0.2 mM IPTG for 16 hours at 16 °C. Cells were harvested at 6000 rpm for 10 minutes and lysed by sonication in buffer containing 25 mM Tris-HCl (pH 8.0), 350 mM NaCl, 30 mM imidazole, 5 mM β-mercaptoethanol, 1 mM PMSF, and 2 mM benzamidine. The supernatant was incubated with Ni-NTA resin (Qiagen) for 30 minutes. Bound protein was eluted using 300 mM imidazole and further purified by Ulp1 tag cleavage followed by ion-exchange. Purified protein was concentrated to ~10–15 mg mL^−1^ and stored at –80 °C. Crystallization of *Bt*ALDH3 was performed by hanging-drop vapor diffusion at 22 °C. Crystals were obtained in two conditions: 190 mM magnesium chloride, 100 mM Tris-HCl (pH 8.5), 21% (w/v) PEG8000, or 200 mM magnesium formate, 100 mM Tris-HCl (pH 8.5), 20% PEG3350

Crystals were harvested, cryoprotected in mother liquor supplemented with glycerol, and flash-frozen in liquid nitrogen for data collection. Diffraction data were collected to 1.4 Å resolution at NSLS II and processed with Mosflm^70^. Crystals of *Bt*ALDH3 (NAD^+^) belonged to space group P1, with unit-cell parameters a = 46.68 Å, b = 61.53 Å, c = 87.95 Å, and α = 94.01°, β = 100.94°, γ = 115.36°. *Bt*ALDH3 (NMN+) crystallized in space group P12 1 1, with a = 93.87 Å, b = 58.98 Å, c = 158.15 Å, and α = γ = 90°, β = 98.41°, γ = 90°. The structure was solved by molecular replacement using an AlphaFold-predicted *Bt*ALDH3 model. 3D structural figures were generated in PyMOL. Final data-collection and refinement statistics are presented in Table S4.

### Molecular Dynamics

Molecular dynamics simulations of apo *Bt*ALDH3 WT, single mutant variants R289A, H290A, R293A, and the corresponding triple mutant were performed as follows: The apo structure was first generated by removing NMN^+^ from its crystal structure, using the first two chains to simulate the functional dimer. The system was solvated using in-house scripts in a rhombic dodecahedron with 10 Å buffer on all sides for a periodic image distance of 120 Å. Simulations utilized the CHARMM36 force field for proteins and the TIP3P model for water. The box was solvated with 100 mM NaCl, and protonation states were assigned at a pH of 7 by ProPKA and visual inspection, resulting in protonation of E63, E210, and E334, δ protonation of H53, H101, and H183, and ϵ protonation of remaining histidines. Simulations of each system at 25 °C were run in triplicate for 250 ns, which considering the two halves of the dimer gave 6 independent trajectories of cofactor pocket dynamics. Simulations were run using the BLaDE module of the CHARMM molecular dynamics package using particle mesh Ewald for the treatment of long-range electrostatics with 6th order interpolation, grid length of 128, and κ = 0.32 Å^−1^. Lennard Jones interactions used force switching with a switch radius of 9 Å and a cutoff of 10 Å. Simulations were analyzed using the MDAnalysis package. For each trajectory, monomers were aligned to an average trajectory structure, and root mean square fluctuation (RMSF) values were computed for Cα atoms across residues 333–336. RMSF values were first averaged across residues for each simulation and then across all six trajectories to quantify residue-level flexibility. Pairwise statistical comparisons between RMSF values of ALDH variants were performed using a two-tailed Welch’s t-test. Trajectories used for structural visualization were subsampled using MDAnalysis, selecting 30 random frames from the trajectory for visualization. Results were visualized with PyMol.

### Multiple Sequence Alignment and Phylogenetic Analysis

Multiple sequence alignments (Fig. 2C) and phylogenetic trees (Fig. 4A) were generated using the EMBL-EBI Clustal Omega webserver tool^71^. The phylogenetic tree shown in Fig. 4A was visualized using the Interactive Tree of Life phylogenetic tree visualizer web tool^72^. Sequence logo plots (Fig. 4B) were generated from the multiple sequence alignments using the WebLogo sequence logo plot generator^73^.

### Synthesis of Non-Nucleotide-Based NRCs

Synthesis of Non-Nucleotide-Based NRCs was carried out using established methods^53^. Briefly, 1 equivalent of nicotinamide was dissolved with 1 equivalent of alkyl bromide in acetonitrile and heated under reflux overnight to yield a white precipitate. The precipitate was filtered and dried under vacuum and analyzed by NMR. The alkyl bromides used for the synthesis of 1-benzylnicotinamide (BNA^+^), 1-(4-carboxy)benzylnicotinamide (BANA^+^), 1-phenethylnicotinamide (P2NA^+^), 1-(3-phenyl)propylnicotinamide (P3NA^+^), 1-(3-(4-methoxyphenyl))propylnicotinamide (P3NA-OMe^+^), and 1-(2-carbamoylmethyl)nicotinamide (AmNA^+^) were (bromomethyl)benzene, 4-(bromomethyl)benzoic acid, (2-bromoethyl)benzene, (3-bromopropyl)benzene, 1-(3-bromopropyl)-4-methoxybenzene, and 2-bromoacetamide, respectively. NMR spectra and detailed reaction conditions are provided in supplementary methods.

### Non-Nucleotide Based Cofactor Conversion Assay

To assess the activity of ALDHs toward non-nucleotide-based cofactors we initially attempted to measure activity in a spectrophotometric assay, measuring either WST-1 formazan in a colorimetric cycling assay or reduced cofactor formation in a specific activity assay (as described above). However, the activity of the tested ALDHs towards these simple synthetic cofactors was too low to accurately quantify using these methods. Instead, we performed a simple conversion assay, measuring the conversion of benzaldehyde to benzoate (Fig. 5). Benzaldehyde was chosen as the substrate as it is utilized by the tested ALDHs, and its product is readily detectable by HPLC. The reactions were carried out as follows: A 20 μL reaction was prepared containing 100 mM Tris-Cl buffer pH 7, 20 mM benzaldehyde, 20 mM cofactor (NAD^+^, BNA^+^, BANA^+^, P2NA^+^, P3NA^+^, P3NA-OMe^+^, AmNA^+^, and “no cofactor”), and 0.25 g L^−1^ purified ALDH. Reactions were initiated by the addition of ALDH to reaction buffer and carried out in PCR tubes at 37 °C for 24 hours. After 24 hours, reactions were arrested by the addition of 40 μL of a 2.5% formic acid, 50% methanol solution. Each sample was then filtered and analyzed by HPLC under the following chromatographic conditions: Separation was achieved on an Agilent 1200 system equipped with a diode array detector (DAD) and Luna Omega Polar C18 column (5 μm, 100 Å, 150 × 4.6 mm, Phenomenex) maintained at 40 °C. The mobile phase consisted of Solvent A: water with 0.1% formic acid and Solvent B: methanol with 0.1% formic acid, run under isocratic conditions at 50% B. The flow rate was 1.0 mL min^−1^, with an injection volume of 5 μL. Total run time was 6 minutes per sample. Detection of benzoate and benzaldehyde was carried out at 240 nm. Concentration of benzoate formed was determined by comparison to authentic standards. All reactions were carried out with *n* = 3 biologically independent replicates.

## AUTHOR CONTRIBUTIONS

S.S., N.H., F.Q., and H.L. designed all experiments. E.L., S.S. and J.B.S performed bioinformatic analysis and sequence similarity network construction. E.L., S.S., W.B.B., J.B.S., and H.L. selected the ALDH candidates. W.B.B. and S.S. constructed ALDH expression vectors. S.S. performed colorimetric assays, 340 nm specific activity assays, and benzaldehyde conversion assays of ALDH candidates. S.S. and V.C.M performed steady-state kinetic characterization of ALDH candidates. N.H. and F.Q. crystallized *Bt*ALDH3. B.S. collected diffraction data. N.H. and J.-K.K. solved and refined the crystal structures. R.L.H and S.S. performed and analyzed molecular dynamics simulations. H.J.C.N. and S.S synthesized non-nucleotide based cofactors. All authors have approved this manuscript for publication.

## ACKNOLEDGEMENTS

S.S., V.M., W.B.B, H.N., and H.L. acknowledge funding from the Advanced Research Projects Agency–Energy EcoSynBio program (Award no. DE-AR0001508), the National Science Foundation (NSF) (Award no. 2328145), the National Institutes of Health (NIH) (Award no. 1R35GM153401-01), Department of Energy (DOE) (Award no. EE0008923) and Sloan Research Fellowship. E.L., and J.B.S. acknowledge funding from Advanced Research Projects Agency–Energy EcoSynBio program (Award no. DE-AR0001508), the National Institute of Environmental Health Sciences (Grant no. P42ES004699), the NIH (Grant no. R01 GM 076324-11), and the NSF (Grant nos. 2328145, 1627539, 1805510, and 1827246). N.H., J.-K.K., and F.Q. acknowledge funding from the NSF (Award no. 2328145). This research used resources at AMX beamline of the National Synchrotron Light Source II, a U.S. Department of Energy (DOE) Office of Science User Facility operated for the DOE Office of Science by Brookhaven National Laboratory under Contract No. DE-SC0012704. The Center for BioMolecular Structure (CBMS) is primarily supported by the National Institutes of Health, National Institute of General Medical Sciences (NIGMS) through a Center Core P30 Grant (P30GM133893), and by the DOE Office of Biological and Environmental Research (KP1605010). The ALS-ENABLE beamlines are supported in part by the National Institutes of Health, National Institute of General Medical Sciences, grant P30 GM124169-01. The Advanced Light Source is a Department of Energy Office of Science User Facility under Contract No. DE-AC02-05CH11231. This work utilized the high-performance computing cluster maintained by the Research Cyberinfrastructure Center (RCIC) at the University of California, Irvine.

The author(s) declare no competing interests.

Supplementary Information is available for this paper.

## REFERENCES

1) Butler, N. D., Anderson, S. R., Dickey, R. M., Nain, P., & Kunjapur, A. M. (2023). Combinatorial gene inactivation of aldehyde dehydrogenases mitigates aldehyde oxidation catalyzed by E. coli resting cells. Metabolic Engineering, 77, 294–305.

2) Montaño López, J., Duran, L., & Avalos, J. L. (2022). Physiological limitations and opportunities in microbial metabolic engineering. Nature Reviews Microbiology, 20(1), 35–48.

3) Bekiaris, P. S., & Klamt, S. (2023). Network-wide thermodynamic constraints shape NAD (P) H cofactor specificity of biochemical reactions. Nature Communications, 14(1), 4660.

4) Rollin, Joseph A., Tsz Kin Tam, and Y-H. Percival Zhang. “New biotechnology paradigm: cell-free biosystems for biomanufacturing.” Green chemistry 15.7 (2013): 1708–1719.

5) Chánique, A. M., & Parra, L. P. (2018). Protein engineering for nicotinamide coenzyme specificity in oxidoreductases: attempts and challenges. Frontiers in microbiology, 9, 194.

6) Weusthuis, R. A., Folch, P. L., Pozo-Rodríguez, A., & Paul, C. E. (2020). Applying non-canonical redox cofactors in fermentation processes. Iscience, 23(9).

7) Knaus, T., Paul, C. E., Levy, C. W., De Vries, S., Mutti, F. G., Hollmann, F., & Scrutton, N. S. (2016). Better than nature: nicotinamide biomimetics that outperform natural coenzymes. Journal of the American Chemical Society, 138(3), 1033–1039.

8) Wang, X., Feng, Y., Guo, X., Wang, Q., Ning, S., Li, Q., Wang, J., Wang, L., & Zhao, Z. K. (2021). Creating enzymes and self-sufficient cells for biosynthesis of the non-natural cofactor nicotinamide cytosine dinucleotide. Nature communications, 12(1), 2116.

9) Campbell, E., Meredith, M., Minteer, S. D., & Banta, S. (2012). Enzymatic biofuel cells utilizing a biomimetic cofactor. Chemical communications, 48(13), 1898–1900.

10) Platt, A. P., Klem, H., Mallinson, S. J., Bomble, Y. J., & Paton, R. S. (2025). Computer-aided design of stability enhanced nicotinamide cofactor biomimetics for cell-free biocatalysis. Green Chemistry.

11) Gu, X., Zhou, J., Wang, H., Zhu, Y., & Ni, Y. (2025). Engineering glucose dehydrogenase from Sulfolobus sulfataricus toward utilizing diverse nicotinamide cofactor biomimetics. Green Synthesis and Catalysis.

12) Zhu, Y., Zhou, J., Gu, X., Wang, H., Han, H., & Ni, Y. (2025). Engineering a newly identified alcohol dehydrogenase from Sphingobium Sp. for efficient utilization of nicotinamide cofactors biomimetics. Bioresources and Bioprocessing, 12(1), 41.

13) Huang, R., Chen, H., Upp, D. M., Lewis, J. C., & Zhang, Y. H. P. J. (2019). A high-throughput method for directed evolution of NAD (P)+-dependent dehydrogenases for the reduction of biomimetic nicotinamide analogues. ACS catalysis, 9(12), 11709–11719.

14) King, E., Maxel, S., & Li, H. (2020). Engineering natural and noncanonical nicotinamide cofactor-dependent enzymes: design principles and technology development. Current opinion in biotechnology, 66, 217–226.

15) Mak, W. S., & Siegel, J. B. (2014). Computational enzyme design: transitioning from catalytic proteins to enzymes. Current opinion in structural biology, 27, 87–94.

16) Black, W. B., Zhang, L., Mak, W. S., Maxel, S., Cui, Y., King, E., Fong, B., Sanchez Martinez, A., Siegel, J.B., & Li, H. (2020). Engineering a nicotinamide mononucleotide redox cofactor system for biocatalysis. Nature chemical biology, 16(1), 87–94.

17) Aspacio, D., Zhang, Y., Cui, Y., Luu, E., King, E., Black, W. B., Perea, S., Zhu, Q., Wu, Y., Luo, R., Siegel, J.B., & Li, H. (2024). Shifting redox reaction equilibria on demand using an orthogonal redox cofactor. Nature Chemical Biology, 20(11), 1535–1546.

18) Richardson, K. N., Black, W. B., & Li, H. (2020). Aldehyde production in crude lysate-and whole cell-based biotransformation using a noncanonical redox cofactor system. ACS catalysis, 10(15), 8898–8903.

19) Zhang, L., King, E., Black, W. B., Heckmann, C. M., Wolder, A., Cui, Y., Nicklen, F., Siegel, J.B., Luo, R., Paul, C.E., & Li, H. (2022). Directed evolution of phosphite dehydrogenase to cycle noncanonical redox cofactors via universal growth selection platform. Nature Communications, 13(1), 5021.

20) Li, Q., Su, H., Meng, D., Qin, Y., Wu, R., Zhu, Z., Sheng, X., You, C., & Job Zhang, Y. H. P. (2025). Stoichiometric Regeneration of Biomimetic Nicotinamide Coenzyme Powered by Biomass Sugars via In Vitro Synthetic Enzymatic Biosystems. ChemSusChem, 18(4), e202401263. Nowak, Claudia, et al. “Characterization of biomimetic cofactors according to stability, redox potentials, and enzymatic conversion by NADH oxidase from Lactobacillus pentosus.” ChemBioChem 18.19 (2017): 1944–1949.

21) Paul, C. E., Tischler, D., Riedel, A., Heine, T., Itoh, N., & Hollmann, F. (2015). Nonenzymatic regeneration of styrene monooxygenase for catalysis. Acs Catalysis, 5(5), 2961–2965.

22) Ryan, J. D., Fish, R. H., & Clark, D. S. (2008). Engineering cytochrome P450 enzymes for improved activity towards biomimetic 1, 4‐NADH cofactors. ChemBioChem, 9(16), 2579–2582.

23) Drenth, J., Yang, G., Paul, C. E., & Fraaije, M. W. (2021). A tailor-made deazaflavin-mediated recycling system for artificial nicotinamide cofactor biomimetics. ACS catalysis, 11(18), 11561–11569.

24) Sicsic, S., Durand, P., Langrene, S., & le Goffic, F. (1984). A new approach for using cofactor dependent enzymes: example of alcohol dehydrogenase. FEBS letters, 176(2), 321–324.

25) Hu, X., Zhang, J., & Rydström, J. (1998). Interactions of reduced and oxidized nicotinamide mononucleotide with wild-type and αD195E mutant proton-pumping nicotinamide nucleotide transhydrogenases from Escherichia coli. Biochimica et Biophysica Acta (BBA)-Bioenergetics, 1367(1-3), 134–138.

26) Shortall, K., Djeghader, A., Magner, E., & Soulimane, T. (2021). Insights into aldehyde dehydrogenase enzymes: a structural perspective. Frontiers in Molecular Biosciences, 8, 659550.

27) Steffler, F., Guterl, J. K., & Sieber, V. (2013). Improvement of thermostable aldehyde dehydrogenase by directed evolution for application in synthetic cascade biomanufacturing. Enzyme and microbial technology, 53(5), 307–314.

28) Park, Y. S., Choi, U. J., Nam, N. H., Choi, S. J., Nasir, A., Lee, S. G., Kim, K.J., Jung, G.Y., Choi, S., Shim, J.Y., & Yoo, T. H. (2017). Engineering an aldehyde dehydrogenase toward its substrates, 3-hydroxypropanal and NAD+, for enhancing the production of 3-hydroxypropionic acid. Scientific Reports, 7(1), 17155.

29) Chen, Z., Wang, C., Wang, R., Li, L., & Du, J. (2025). Engineering of an Aldehyde Dehydrogenase from Neurospora crassa for Crocetin Production. Journal of Agricultural and Food Chemistry.

30) Kanter, J. P., Honold, P. J., Lüke, D., Heiles, S., Spengler, B., Fraatz, M. A., Harms, C., Ley, J.P., Zorn, H., & Hammer, A. K. (2022). An enzymatic tandem reaction to produce odor-active fatty aldehydes. Applied microbiology and biotechnology, 106(18), 6095–6107.

31) Zhou, J., Tian, X., Yang, Q., Zhang, Z., Chen, C., Cui, Z., Ji, Y., Schwaneberg, U., Chen, B., & Tan, T. (2022). Three multi-enzyme cascade pathways for conversion of C1 to C2/C4 compounds. Chem Catalysis, 2(10), 2675–2690.

32) Bar-Even, A., Noor, E., Savir, Y., Liebermeister, W., Davidi, D., Tawfik, D. S., & Milo, R. (2011). The moderately efficient enzyme: evolutionary and physicochemical trends shaping enzyme parameters. Biochemistry, 50(21), 4402–4410.

33) Black, W. B., Perea, S., & Li, H. (2023). Design, construction, and application of noncanonical redox cofactor infrastructures. Current Opinion in Biotechnology, 84, 103019.

34) Jackson, B., Brocker, C., Thompson, D. C., Black, W., Vasiliou, K., Nebert, D. W., & Vasiliou, V. (2011). Update on the aldehyde dehydrogenase gene (ALDH) superfamily. Human genomics, 5(4), 283.

35) Suzek, B. E., Wang, Y., Huang, H., McGarvey, P. B., Wu, C. H., & UniProt Consortium. (2015). UniRef clusters: a comprehensive and scalable alternative for improving sequence similarity searches. Bioinformatics, 31(6), 926–932.

36) Zallot, R., Oberg, N., & Gerlt, J. A. (2019). The EFI web resource for genomic enzymology tools: leveraging protein, genome, and metagenome databases to discover novel enzymes and metabolic pathways. Biochemistry, 58(41), 4169–4182.

37) Aspacio, D., Luu, E., Worakaensai, S., Cui, Y., Maxel, S., King, E., Hagerty, R., Chu, A., Minn, D., Siegel, J.B., & Li, H. (2024). Engineering Escherichia coli Pyruvate Metabolism to Generate Noncanonical Reducing Power. ACS Catalysis, 14(13), 9776–9784.

38) King, E., Cui, Y., Aspacio, D., Nicklen, F., Zhang, L., Maxel, S., Luo, R., Siegel, J.B., Aitchison, E., & Li, H. (2022). Engineering Embden–Meyerhof–Parnas glycolysis to generate noncanonical reducing power. ACS catalysis, 12(14), 8582–8592.

39) Chen, Y. L., Chou, Y. H., Hsieh, C. L., Chiou, S. J., Wang, T. P., & Hwang, C. C. (2022). Rational engineering of 3α-hydroxysteroid dehydrogenase/carbonyl reductase for a biomimetic nicotinamide mononucleotide cofactor. Catalysts, 12(10), 1094.

40) Kim, J. Y., King, E., Cui, Y., Zhang, L., Siegel, J. B., & Li, H. (2025). Engineering the key central metabolic enzyme to utilize a non-canonical redox cofactor. bioRxiv, 2025-05.

41) Liu, Z. J., Sun, Y. J., Rose, J., Chung, Y. J., Hsiao, C. D., Chang, W. R., Chang, W.R., Kuo, I., Perozich, J., Lindahl, R., Hempel, J., & Wang, B. C. (1997). The first structure of an aldehyde dehydrogenase reveals novel interactions between NAD and the Rossmann fold. Nature structural biology, 4(4), 317–326.

42) Khanna, M., Chen, C. H., Kimble-Hill, A., Parajuli, B., Perez-Miller, S., Baskaran, S., Kim, J., Dria, K., Vasiliou, V., Mochly-Rosen, D., & Hurley, T. D. (2011). Discovery of a novel class of covalent inhibitor for aldehyde dehydrogenases. Journal of Biological Chemistry, 286(50), 43486–43494.

43) Zahniser, M. P., Prasad, S., Kneen, M. M., Kreinbring, C. A., Petsko, G. A., Ringe, D., & McLeish, M. J. (2017). Structure and mechanism of benzaldehyde dehydrogenase from Pseudomonas putida ATCC 12633, a member of the Class 3 aldehyde dehydrogenase superfamily. Protein engineering, design and selection, 30(3), 273–280.

44) Perez-Miller, S. J., & Hurley, T. D. (2003). Coenzyme isomerization is integral to catalysis in aldehyde dehydrogenase. Biochemistry, 42(23), 7100–7109.

45) Huo, L., Davis, I., Liu, F., Andi, B., Esaki, S., Iwaki, H., Hasegawa, Y., Orville, A.M., & Liu, A. (2015). Crystallographic and spectroscopic snapshots reveal a dehydrogenase in action. Nature communications, 6(1), 5935.

46) d’Ambrosio, K., Pailot, A., Talfournier, F., Didierjean, C., Benedetti, E., Aubry, A., Branlant, G., & Corbier, C. (2006). The first crystal structure of a thioacylenzyme intermediate in the ALDH family: new coenzyme conformation and relevance to catalysis. Biochemistry, 45(9), 2978–2986.

47) Boehr, David D., et al. “The dynamic energy landscape of dihydrofolate reductase catalysis.” science 313.5793 (2006): 1638–1642.

48) Zou, Y., Zhang, H., Brunzelle, J. S., Johannes, T. W., Woodyer, R., Hung, J. E., Nair, N., Van Der Donk, W.A., Zhao, H., & Nair, S. K. (2012). Crystal structures of phosphite dehydrogenase provide insights into nicotinamide cofactor regeneration. Biochemistry, 51(21), 4263–4270.

49) Cristobal, J. R., Nagorski, R. W., & Richard, J. P. (2023). Utilization of cofactor binding energy for enzyme catalysis: Formate dehydrogenase-catalyzed reactions of the whole NAD cofactor and cofactor pieces. Biochemistry, 62(15), 2314–2324.

50) Colonna-Cesari, F., Perahia, D., Karplus, M., Eklund, H., Brädén, C. I., & Tapia, O. (1986). Interdomain motion in liver alcohol dehydrogenase. Structural and energetic analysis of the hinge bending mode. Journal of Biological Chemistry, 261(32), 15273–15280.

51) Price, A. C., Zhang, Y. M., Rock, C. O., & White, S. W. (2004). Cofactor-induced conformational rearrangements establish a catalytically competent active site and a proton relay conduit in FabG. Structure, 12(3), 417–428.

52) Skarżyński, T., & Wonacott, A. J. (1988). Coenzyme-induced conformational changes in glyceraldehyde-3-phosphate dehydrogenase from Bacillus stearothermophilus. Journal of molecular biology, 203(4), 1097–1118.

53) King, E., Maxel, S., Zhang, Y., Kenney, K. C., Cui, Y., Luu, E., Siegel, J.B., Weiss, G.A., Luo, R., & Li, H. (2022). Orthogonal glycolytic pathway enables directed evolution of noncanonical cofactor oxidase. Nature Communications, 13(1), 7282.

54) Henzler-Wildman, K. A., Thai, V., Lei, M., Ott, M., Wolf-Watz, M., Fenn, T., Pozharski, E., Wilson, M.A., Petsko, G.A., Karplus, M., & Kern, D. (2007). Intrinsic motions along an enzymatic reaction trajectory. Nature, 450(7171), 838–844.

55) Schwartz, S. D. (2023). Protein dynamics and enzymatic catalysis. The Journal of Physical Chemistry B, 127(12), 2649–2660.

56) Boehr, D. D., Nussinov, R., & Wright, P. E. (2009). The role of dynamic conformational ensembles in biomolecular recognition. Nature chemical biology, 5(11), 789–796.

57) Zhou, J., Gu, X., Chen, S., Zhu, Y., & Ni, Y. (2025). Biooxidation of Aromatic Aldehydes by Aldehyde Dehydrogenase from Sphingobium sp. Using Synthetic Nicotinamide Cofactor Biomimetics. Green Chemical Engineering.

58) Lauko, A., Pellock, S. J., Sumida, K. H., Anishchenko, I., Juergens, D., Ahern, W., Jeung, J., Shida, A.F., Hunt, A., Kalvet, I., & Baker, D. (2025). Computational design of serine hydrolases. Science, 388(6744), eadu2454.

59) Guo, A. B., Akpinaroglu, D., Stephens, C. A., Grabe, M., Smith, C. A., Kelly, M. J., & Kortemme, T. (2025). Deep learning–guided design of dynamic proteins. Science, 388(6749), eadr7094.

60) Otten, R., Pádua, R. A., Bunzel, H. A., Nguyen, V., Pitsawong, W., Patterson, M., Sui, S., Perry, S.L., Cohen, A.E., Hilvert, D., & Kern, D. (2020). How directed evolution reshapes the energy landscape in an enzyme to boost catalysis. Science, 370(6523), 1442–1446.

61) Macheroux, P., Kappes, B., & Ealick, S. E. (2011). Flavogenomics–a genomic and structural view of flavin‐dependent proteins. The FEBS journal, 278(15), 2625–2634.

62) Heinonen, J. K. (2012). Biological role of inorganic pyrophosphate. Springer Science & Business Media.

63) Igamberdiev, A. U., & Kleczkowski, L. A. (2021). Pyrophosphate as an alternative energy currency in plants. Biochemical Journal, 478(8), 1515–1524.

64) Baykov, A. A., Malinen, A. M., Luoto, H. H., & Lahti, R. (2013). Pyrophosphate-fueled Na+ and H+ transport in prokaryotes. Microbiology and Molecular Biology Reviews, 77(2), 267–276.

65) Lindahl, R., & Petersen, D. R. (1991). Lipid aldehyde oxidation as a physiological role for class 3 aldehyde dehydrogenases. Biochemical pharmacology, 41(11), 1583–1587.

66) Liu, S., & Zhang, W. (2023). NAD+ metabolism and eye diseases: current status and future directions. Molecular Biology Reports, 50(10), 8653–8663.

67) Imai, S. I. (2025). NAD World 3.0: the importance of the NMN transporter and eNAMPT in mammalian aging and longevity control. npj Aging, 11(1), 4.

68) Sorci, L., Martynowski, D., Rodionov, D. A., Eyobo, Y., Zogaj, X., Klose, K. E., K.E., Nikolaev, E.V., Magni, G., Zhang, H., & Osterman, A. L. (2009). Nicotinamide mononucleotide synthetase is the key enzyme for an alternative route of NAD biosynthesis in Francisella tularensis. Proceedings of the National Academy of Sciences, 106(9), 3083–3088.

69) Gazzaniga, F., Stebbins, R., Chang, S. Z., McPeek, M. A., & Brenner, C. (2009). Microbial NAD metabolism: Lessons from comparative genomics. Microbiology and Molecular Biology Reviews, 73(3), 529–541. 10.1128/mmbr.00042-08

70) Battye, T. G. G., Kontogiannis, L., Johnson, O., Powell, H. R., & Leslie, A. G. (2011). iMOSFLM: a new graphical interface for diffraction-image processing with MOSFLM. Biological crystallography, 67(4), 271–281.

71) Madeira, F., Pearce, M., Tivey, A. R., Basutkar, P., Lee, J., Edbali, O., Madhusoodanan, N., Kolesnikov, A., & Lopez, R. (2022). Search and sequence analysis tools services from EMBL-EBI in 2022. Nucleic acids research, 50(W1), W276–W279.

72) Letunic, I., & Bork, P. (2021). Interactive Tree Of Life (iTOL) v5: an online tool for phylogenetic tree display and annotation. Nucleic acids research, 49(W1), W293–W296.

73) Crooks, G. E., Hon, G., Chandonia, J. M., & Brenner, S. E. (2004). WebLogo: a sequence logo generator. Genome research, 14(6), 1188–1190.

