## Supplementary Information for "A Sequence Motif Enables Widespread Use of Non-Canonical Redox Cofactors in Natural Enzymes"

[1]Department of Chemical and Biomolecular Engineering, University of California, Irvine, Irvine, CA, USA
[2]UCI Center for Synthetic Biology, University of California, Irvine, Irvine, CA, USA
[3]Department of Biological Chemistry, University of California, Irvine, Irvine, CA, USA
[4]Genome Center, University of California, Davis, Davis, CA, USA
[5]Department of Chemistry, University of California, Davis, Davis, CA, USA
[6]Department of Biochemistry and Molecular Medicine, University of California, Davis, Davis, CA, USA
[7]Department of Pharmaceutical Sciences, University of California, Irvine, Irvine, CA, USA
[8]Department of Chemistry, University of California, Irvine, Irvine, CA, USA
[9]Molecular Biophysics and Integrated Bioimaging, Berkeley Center for Structural Biology, Lawrence Berkeley National Laboratory, Berkeley, CA, USA
[10]These authors contributed equally.
*To whom correspondence should be addressed

**CORRESPONDING AUTHORS**

Han Li - Department of Chemical and Biomolecular Engineering, University of California, Irvine, Irvine, CA, USA; UCI Center for Synthetic Biology, University of California, Irvine, Irvine, CA, USA; Department of Biological Chemistry, University of California, Irvine, Irvine, CA, USA


Feng Qiao - Department of Biological Chemistry, University of California, Irvine, Irvine, CA, USA


**SUPPLEMENTARY DATA**

Figure S1: Complete Specific Activity Data of *Bt*ALDH3, *Pb*ALDH, and Related Homologs with $NAD^+$ and $NMN^+$

Figure S2: Activity of *Sp*ALDH2 with Simple Synthetic Cofactors

Table S1: Plasmid and Strain List

Table S2: Amino Acid Sequences of Enzymes Used

Table S3: Colorimetric Screening Data

Table S4: Data Collection and Refinement Statistics

Table S5**:** Apparent Kinetic Parameters of *Hs*ALDH3 and *Pp*MdlD with $NAD^+$

**SUPPLEMENTARY METHODS**

Synthesis and Characterization of 1-benzylnicotinamide bromide

Synthesis and Characterization of 1-(4-carboxy)benzylnicotinamide bromide

Synthesis and Characterization of 1-phenylethylnicotinamide bromide

Synthesis and Characterization of 1-(3-phenyl)propylnicotinamide bromide

Synthesis and Characterization of 1-(3-(4-methoxyphenyl))propylnicotinamide bromide

Synthesis and Characterization of 1-(2-carbamoylmethyl)nicotinamide bromide

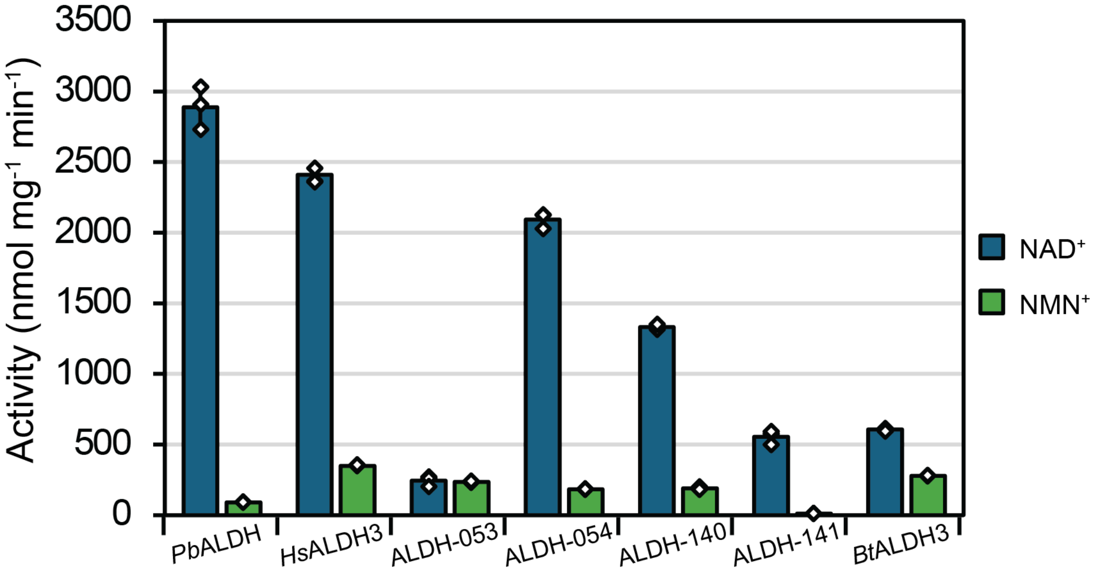


**Figure S1: Complete specific activity data of *Bt*ALDH3, *Pb*ALDH, and related ALDHs with NAD$^+$ and NMN$^+$.** ALDHs were tested under standard specific activity assay conditions as described in methods. All ALDH's were tested with hexanal except for ALDH-054, which was tested with acetaldehyde. All specific activity data were generated by monitoring the formation of reduced cofactor (NADH or NMNH) at 340 nm and was carried out for $n$ = 3 independent biological replicates. Error bars correspond to one standard deviation.

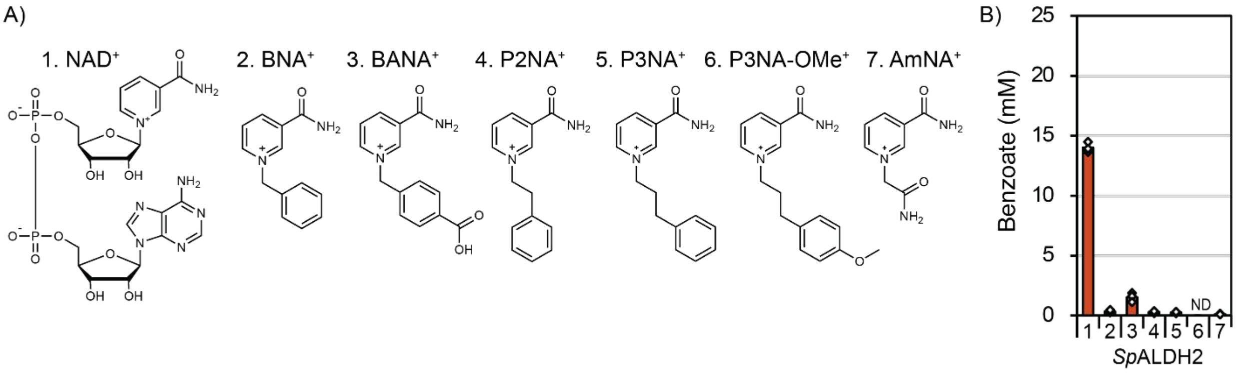


**Figure S2: Activity of *Sp*ALDH2 with Simple Synthetic Cofactors.** *Sp*ALDH utilizes simple synthetic, non-nucleotide based cofactors under the conditions described in methods. All reactions were carried out with $n$ = 3 biologically independent replicates. "ND" represents no benzoate detected.

**Table S1:** Plasmid and Strain List

| Strains | Description | Note |
|---|---|---|
| XL-1 Blue | *E. coli* cloning strain | Stratagene |
| BL21 | *E. coli* protein expression strain | Invitrogen |
| **Plasmids** | **Description** | **Note** |
| pSS687 | T7:: *Bt*ALDH3, pBR322 ori, KanR, N-terminal 10xHis-Smt3 tag | Used to express *Bt*ALDH3 for crystallographic studies |
| pQElac | PLlacO1:: empty, ColE1 ori, AmpR, N-terminal 6x His-tag | Empty vector used in recombinant protein expression and purification of ALDHs[1] |
| pW112 | PLlacO1:: Geobacillus sp. (strain Y412MC52) azoR, ColE1 ori, AmpR, N-terminal 6x His-tag | Referred to as pQE_*Gs*DI in this work. Uniprot: A0A0E0THL1. |
| ALDH-### | PLlacO1:: ALDH-###, ColE1 ori, AmpR, N-terminal 6x His-tag | ALDHs used in this work were all cloned into pQE. Refer to Table S2 for full sequence. |

**Table S2:** Amino Acid Sequences of Enzymes Used

| Plasmid # | UniProt/UniParc ID | Sequence |
|---|---|---|
| pW112 | A0A0E0THL1 | MTKVLYITAHPHDDTQSYSMAVGKAFIETYKQVHPDHEVIHLDL<br>YKEYIPEIDVDVFSGWGKLRSGKSFEELSDEEKAKVGRMNELCE<br>QFISADKYVFVTPMWNFSFPPVLKAYIDAVAVAGKTFKYTEQGP<br>VGLLTDKKALHIQARGGFYSEGPAAEMEMGHRYLSVIMQFFGV<br>PSFEGLFVEGHAAMPEKAEEIKANAIARAKDLAHTF* |
| ALDH-009 | A0A0D6KE96 | MTTQLTPPQSQTTIRTYQNFINGKYISPLSGVNYERKSPLTGETIV<br>QIPWSNQADTDVAIQAARQVFDDGTWSTSHARVRHDILRKTAEL<br>LTTKTSEIAAVICQEVGRPIGMCIGEVQMTAQVFDYFAALTLNLQ<br>GESTTQYDRNAIGLTVHEPVGVVGIITPWNFPLLLVAWKIAPAIAA<br>GCTMVVKPSEFTPSTAFMLAEILSEAGLPDGVINIVTGDGPVVGE<br>HLVESPLVDKIAFTGSTAVGRRIMAKGAPTLKRMSLELGGKSPNI<br>VFGDADLSQAIPGALFGIYINSGQVCQAGSRLLLHESIKDVFIEQF<br>LAATQTFQIGNPTDNTTMMGPVINEIQFERIQNYIQLGEKEGAKL<br>LIGGSGRYLVPGFEQGLFIKPTVFDHVTNEMAIAQEEIFGPVLSIM<br>TFKDKAEALQIANQTMYGLTAAIWTKNLDTAFKMAKGIRAGTV<br>WVNSYHTSGLEPTMPYGGYKQSGIGREVGKNGLEEYLETKAIHI<br>KLA* |
| ALDH-010 | L8N0N6 (*Pb*ALDH) | METLPINNLDIPATIAKQRIFFDGNKTKDYEFRVSQLKKLAQLIKE<br>NEKLILDAVYADLRKPTIEIFGSEILIALSEIRYVIKHLKAWMKPQK<br>VGTPLNLFPSSSYIYTEPLGVVLIVAPWNYPFSLNIQPLIGAIAAG<br>NCAILKPSEYAPHTSNAIAKIINEHFDPNFITVIEGGLEINQALLAE<br>KFDHIFFTGSTAIGKIVMEAAAKHLTPVTLELGGKSPCIVDEECDL<br>ETTAKRIIWGKFYNAGQTCVAPDYLLVSKSIKPVLIEKLLGYVKT<br>FFGENPQQSPDFARIVNDRQFDRLVSLLNEGKILIGGQTDKSDRYI<br>APTIIDGISIHAKIMGEEIFGPILPVLEYDQLSEAIALIKSQSQPLAL<br>YLFSNNKQKQEKILQEISFGGGCFNDTILHLANLELPFGGVGNSG<br>MGSYHGKATFDRFSHRKSVLKNSFRFDLKLRYPPYRVSIDTLKK<br>FIN* |
| ALDH-011 | A0A2V7VB71 | LKAMAGERLDIPLVIGGKEVRTGDKAKAVMPHDHRHVLGDWH<br>KASREHVAQAIDAAAKAHGEWSRWPWEDRAAVFLKAADLLAT<br>TWRATLNAATMLGQSKTVFQAEIDSACELVDFWRFNPAYAQELY<br>AEQPLSSAGMWNQSEYRPLEGFVYAITPFNFTAIGGNLPTSPALM<br>GNTVVWKPASAAIPSGYWIMKLLEAAGLPPGVVNFVPGDAVTV<br>SDTVLTHRDLAGVHFTGSTEVFNSMWKTIGGSMSRYRSYPRIVG<br>ETGGKDFIVAHPSAEPQALAVAIARGGFEYQGQKCSAASRVYVP<br>RSLWSDVRDRTVAIIKEIRVGDVTDFRNFMGAVIDKKAFDKISEYI<br>GDARKNATIVAGGGANGETGYFIEPTLVEAREPGYRLLCEEIFGP<br>VVTVYVYPDEKWEETLAVVDQTSAYALTGAVFATDRGAVRQAA<br>SALRHAAGNFYVNDKPTGAVVGQQPFGGSRGSGTNDKAGSKLN<br>LVRWVSARSIKETFNPPRDYRYPFMAEE* |
| ALDH-012 | UPI000407F0B2 | MNIPLEPLAATTDRRIEAIALGRTYTTQRRAIVHDLLSIPVAEMTV<br>APPIYIHKAISSMREAPSLSLAEIHAAMARAADSYQYDTIAGLSP<br>DEYSRLLQRTTGLPETVTQNALATVADALRNMPDIINAGRPQGA<br>RWSWDDADALAGYSLFSRKGDVFAVLAAGNGPGIHALWPQAVA |

| | | |
|---|---|---|
| | | LGYRTLVRPSTREPFTAQRVICAMVQAGLEHYVALIPTDYRGADE LVASADLALAYGGQDIVDKYRHHPQVRVQGPGRAKILIGADVC MEEAVSLVATSMTDLGGAACVSASAVMVEGDVSDFCRRLRQTL QQLPEKVLPRASQQTVDWLNSVIDTPIEASLMSEGYLLRPVVTE VAEPDDPRIHRELPFPCVTVAPYHPTRSAQILSGSLVVTVFSRQPQ LLKSIIADASISNVYVGDIPTTWMSPLVPHDAYLSDFLMCNRGFRI AASWIEAESKGEKL* |
| ALDH-013 | A0A2Z4LU87 | MFKFNISGRNFIGYNRSSKGDTTFQAKNPSTGEGLITSYYEATLD EVNQAIELAEKAFTGYREKTGQEKASFLEAIAEELAQLEEGLVG LCVQETGLPEGRLKGELGRTMGQLRLFASVLREGSWVDARIDFA VPDRKPSPRPDIRYMQKSLGPVGIFGASNFPLAFSVAGGDTASAL AAGCSIVVKAHPSHPGTSELVAMAINEAAKKSNMPDGIFSMLHG VSNTVGEAIVQHPLIKAIGFTGSFKGGKALYDKAVRRKEPIPVYA EMGSVNPIFILSNALKEQYKTIAKGLSDSVQMGVGQFCTNPGITI VPNINETSLFKEELNNCISNSESATMLSASIQEGYETGLKRLKSNE IITSLSKGEQKEGHNQGVPEILSVSAKDFLSENTLEEEVFGPSTLL VHAQDGEEMLKIAKSLHGHLTATIHGTEEDLQANVDLLKILERK VGRLLINGFPTGVEVCHSMVHGGPFPATTDSRMTSVGTAAITRFT RPICFQNFPDVLLPDELKDGNPLKILRMENGKYMKAD* |
| ALDH-014 | A0A0S8BPB2 | MQGSAAIPEAPHRIEPTSQSKLDDALAILDDHKQEWAALDLEERI ELLEQMREGVVEVAEDWVRASVEAKGMTFGTAEEGEEWMEGP ATTLRNIMLLIAALRDIEEYGVPQLPKPAFTRPDGQVVAPVLPAST WDKLLFQGFTAEVWMQPGVTLENMSESQAAFYRDKAPKGRVA LVLGAGNVASIGPMDALYKLFVEGQVVILKMNPVNEYLGPFIDH AFAALRERGFFRVVYGGAAEGDYLCRHERVEEIHITGSDKTHDA IVFGFGEEGSRRKAEAEPRNTKRLTCELGNVSPVIIVPGPWSQKD LDFHGVNLATSVVNNAGFNCSATRVIIQHEQWGKREALLASLRK AFQQAEDRRPYYPGSEERQQLFLEHHPHAEEFGTKGEGHVPWT LIHHLDPSDPDEICFNTESWCGQTSEVALPADSVAEYIDRAVEFCN ERVWGTLNCSIFVHPKSMKDPQIAAAVDRAIASLRYGAIAVNHW AGLNFALVTPTWGAFPGHTTEDIRSGRGVVHNTYMFDRPQKSV VRGPFRVFPKPAWFIDHKTAAEVGRKMTYFNADPSLARLPGLIW SSLFG* |
| ALDH-015 | UPI000E14A9E8 | MTPVINPAEPKDIVGYVREATHAEVEQALQNAANNAPIWFATPP QERAAILHRAAVLMEGQMQQLIGILVREAGKTFSNAIAEVREAV DFLHYYAGQVRDDFDNETHRPLGPVVCISPWNFPLAIFTGQIAA ALAAGNSVLAKPAEQTPLIAAQGIAILLEAGVPPGVVQLLPGQGE TVGAQLTSDERVRGVMFTGSTEVATLLQRNIATRLDAQGRPIPLI AETGGMNAMIVDSSALTEQVVVDVLASAFDSAGQRCSALRVLC LQDDVADHTLKMLRGAMAECRMGNPGRLTTDIGPVIDKEAKTN IERHIQAMRAKGRPVFQAVRDNSDDAREWQTGTFIAPTLIELESF DELQKEVFGPVLHVVRYTRNNLGSLIEQINASGYGLTLGVHTRID ETIAQVTGSAHVGNLYVNRNMVGAVVGVQPFGGEGLSGTGPKA GGPLYLYRLLANRPENALGITLARQDADYPVDAQLKAALVQPLE ALREWATDRPSLHALCQQFGELAQAGTQRLLPGPTGERNTWTL |

| | | |
|---|---|---|
| | | LPRERVLCIADDEQDALVQVAAVASVGSLILWPDDAFHRELAKR<br>LPAAVSGRIQFAKADNHHRAAV* |
| ALDH-016 | A0A7Y1V182 | MPEVFQTISPVDGRVYVERSYAQLQDIDHTLNQAQQIQMEWQAT<br>SIDERAQICRKAVHYLVAHSNKLAQELTWQMGRPIRYTPNEILGG<br>LQERAIHMIDIAASALAEESLEDGNQFKKVIKHGALGTILVLAPW<br>NYPYLTSINAIIPALMAGNTVILKHSQQTPLCAESYADAFNNAGL<br>PAGVFQYLHMTHTQVAKVVVDSRIDFVSFTGSVEGGYAIQEAVG<br>KRFMRTGLELGGKDPAYVRADADLSYCSEQIADGAFFNSGQSCC<br>GVERIYVHTEVYDQFLEAFLSATTQLNLDDPTKADTTLGPMIKPT<br>AAAFVTDQIDAALAMGARPLVDVRSFPNHEVKRGYMAPQVLV<br>DVDHRMSIMQDETFGPAVGIMKVKDDAEAVHLMNDSRYGLTAS<br>IWTQDLERGEVLGKSIQTGTLFINRCDYLDPSLAWSGVKDSGIGI<br>SLSHLGYLQVTRPKSFHIKK* |
| ALDH-017 | A0A517R638 | MMLWKPRPLLPDMIFSDRTVAATVSMDRPNMSTITAQRPNIDRR<br>SHSDDPDEVAFVSEALTQARSSQSAWAEQPLREKLKIVRRFREQI<br>AARPDDFTYAVELPQRRNRAETLASELLPLADACQFLESEAPRLL<br>EPKKLSKRGRPGWLKGVIAEEHREPFGVVLIIATWNYPLLLPGV<br>QMLQALVAGNAVLLKPGKGSHAAAVALREAVVACGLDPNLVTV<br>LSEATSAAQTAITPPEGTSGADKVILTGAAATGRKVLAQTAEHLT<br>PAAMELSGCDAVYVRGDADLDLVVDCLALGMTFNGSATCIAPR<br>RVLVHNSICDELADRLSARFAELPSAAIEPTLAGRLSRLVDEACD<br>DGATKLAGQVEGHVVAPILLRDASPEMALLQADIFAPLLSIVRVT<br>GDEQALEFDRRCPYALGATIFSTDETAARSLAAQINAGCVVINDF<br>LIPTADPRIAFGGRGESGFGVTRGGQGLIEMTQPKTVVVQRASW<br>RPHLTPPDETYEQFFKNYIASAHAKSPWARAKAGFAFLSEAVKR<br>QREQR* |
| ALDH-019 | A0A346XWA7 | MERLEKLTAGMPLLVGGDRFTTVPTDIADAFEPGDAVLVADTGE<br>VLHVPDAERRAATVAVDAAVAAFDLMGTATDEQVSAFYDTFAN<br>ALADDTIFVAIAEANAADVEAARQKGRSTTRLELTTRMRNDMV<br>EGLRGWRDMPGGRGEVVETVTHDGWRVEQVRDRLGVVGFVFE<br>GRPNVFADACGVLRSGNTVVFRIGSDALGTARAIVTHALRPALA<br>AAGLHEGSAALVDSPSRAAGYALFSDPRLSLAVARGSGRAVSML<br>GGIARKAGIPASLHGTGGAWIVAGPTADADDFGEAVTRSLDRKV<br>CNTLNTCAITADRADELVPVFLDALRAAGTARGVEPKLHVSADS<br>VEHVPSDWFDRQVPIARAEGEVKEAQAERIAADELGREWEWED<br>SPEVTLVVVDSIDDAVQRFNAQAPRFVASLLATDPAEHDRFFATID<br>APFVGNGLTRWVDGQYALGKPELGLSNWETGRLFARGGILAGD<br>SVHTVRTRAVLDRPDIPR* |
| ALDH-020 | A0A2E3KN37 | MMGGTGGRTRTDTSAVVYHRRPKPTAPTMTSHSRQLELLRADR<br>SRWAGRFRHLILKGTEEFVRLADSELGKPRHETITAELLPLIASCR<br>WHQRQARRILKTRRLKGRPIWLFGQQHRIQHVPLGTVGIIATWN<br>YPIQLLGIQMLQAALAGNDLIIKPSEHAPQCQAFLIELAHKAGLD<br>ERALRSLPATREAGAAMIERESLDHLVFTGSTRVGRLVAEACASR<br>LIPSTLELSGRDSALVLEDADVQLAARSIWTAVTSNAGQTCMAP<br>RRVLVHQDRYQAFCAALTPLAEKAIPRRLTLPEDAARIDSQVDEA<br>VRAGGRAIPHTEAQSDASAFLPRVVLDCPDGTPLMDGDHFGPAL |

| | | |
|---|---|---|
| | | AVHACASLNEMLEHHQGVGQYLATSIWTGNPTAARGLAAELRS<br>ATVTINDVLIPTAHPGASISGHGPSGWGTSRGAAGLLSMSRPVHV<br>TTTPRKMRMPTDVPTDAALAKLEKLIGVRRPSVDDDSKVTSNSP<br>QPHSGKSS* |
| ALDH-021 | UPI000A40ADAD | MLGASLDKDQPDNPINYFHRIPAHAKRLTTYRTMMNIFTACHPE<br>DLDTAYHYLLKHEKTRHDIINDDTFATVIEKAFAYYKDVIAPAYR<br>PRLLDTQSAALLHSFSAQLKRLDVKNKFITCLLLNPIVDSVGFRC<br>QRDAYKTIYACLFGDEHGPRLAEYINFLGVQAFCEKLDSRIQQA<br>AHDREYDPMKEDTHASQEALQMSDAANHTAKSETLNTSDQAPE<br>AIDFDEIKARAEAYALHLRSHSESIAESLSGFECYNVAVDEIERCIE<br>FLENIELNRSFFERRVNCVTSYLPLNQPIYATTCFGIIPSLLARSAW<br>LRPPTAMHPHYKKLLNPLKLDHFFPNLHVSFADKDTFVSQTAAIS<br>DAVIFTGTPENAAKVRKSYLKRTLFILNGAGHNPLVVAADACITT<br>AVESALRVVLYNQGQDCAGPNSIMVHVEAYPEFIRQLRKELTRC<br>EGLVGDYKCKKNIVGPNSDPDHTLKVMKMFRDCREHCTYGGEI<br>NPVSGLIRPTIFERPLSLGGNYKEFFAPVFFVQLYSDDAELASYFE<br>HPLYSPNSMYISLFGSSDYISGLIEQGKHDPCTVLSNTDLHIVEKG<br>YAPYGGQGVAASCLYVNGVRIAKPTLPQRDIHEHLVAHPHH* |
| ALDH-022 | A0A139NAP6 | MTQTAQEAFAQLDPQAWTKVSIEERLQILAEIQANMRQYGAELG<br>QAEMAMKNRLTGADLYSQTDGMLQTLVAVGNVINASTFIYQTL<br>AETGKMPEAKSIRDLGDGTFEVEVFPTAPVDQMTAATQHGYLRL<br>VGQPKQVSPLDKEAGIIAVSGAGNYSSSIETIKAIFFDNKTVIHKA<br>HRLNEATDKVWEKIFAPLVERKALSFAGVDYSRDLIQLEGLDAIY<br>FTGSTAVAKNIMASTDTPLVSECGGNNPAIIVPGDRPWTAEEIKNQ<br>AELIVSISKGNGGAACGRPQTFITSKQWVQREEFLDAIRQAAQST<br>FAVGTYYPKSADVREAFLAAHPQAEIIKPEGGQYPNTDFLFIPDM<br>DKSAYGVTHEAFCQIMGEVALDVPATAEAFLPAATAFANDELLGT<br>LGCMILIDDETRANHEASFQTALSELNYGGITVNTTPPMVWFNA<br>YLTWGGCKETKENFVSGIGNFGNALNFEQVEKSILVEQFAATGFL<br>YNDRQATDAMNQQVINFTLGNME* |
| ALDH-023 | A0A1X6ZLY8 | MTATPDPALEAALDAFGPAEPAPDRAARKAHLEALEREMRSHAE<br>AAAEAVAGDFGARPRPETLLTEVAMVIGAAQHARRHLRRWMRP<br>ERVVLPPHLWPSTARVDRVPLGRVGIIGPWNYPVQLALVPLVAAI<br>AGGNRAILHPSEHTPRSAALIARIVERAIPDRARVLTGGADQARA<br>LAAAPLDGLFFTGSTATGRHIMAAAAQNLVPVVLELGGKSPAILR<br>HDADIDAAARSIMAGKLLNAGQTCVAPDYAMVPREMLDRFVAA<br>LKTATEALYPDPAGPDYAAIARASDRDRLAALLDGLDPVPLMAR<br>PPAPPRMGAVAVIDPDPDHPLMREEIFGPILPVIPYDAPHEPRDFVA<br>ARPCPLALYVYGRDLAAARSEAEAIPAGGAVINEAVLHVGVQQL<br>PFGGAGASGLGAYHGAEGFRAFTRPRSTMIARPSLARLVRPPYG<br>RNVERILKSLIG* |
| ALDH-024 | A0A060QGV9 | MSDITTSFTRNSHGPAESLEASASLHDECEDKSSSTSSQEVCRDL<br>KAMQPRWAAVPLRARLRVLRRFNRLMLLNAPSLIRLIADHASLD<br>VMTAEILPLMAASRFLCREASSMLKETRLGWLGRPLWLQGVMA<br>SVVRKPLGAVLILAPGNYPLMLASIQTLQALVAGNSVALKPAPGR<br>TAVLRRFVALLEQAGLPKGVVQLVGEDSGQQAVSSGYDLIILTGS |

| | | |
|---|---|---|
| | | AETGRKVALAAAETLTPTIMELSGADPVFVLPDADLALVARALH FGQTLKGGHTCIAPRRIFINEVQKRPLQRELYRVFGGEGDNPAPD PSSKLAQLICSAKAAGGEVVVCGKTQVIFLNAHQARLADIDLFA PWFAVITTHSVEEAICLEGAATHALGASIFGNEREALAIAKRIPAG TITINDIIVPSADPRLPFGGAHRSGFGVTRGREGLLALTRPVSISTR KRGAFHLLPRLRQWHGR* |
| ALDH-025 | UPI0001E31496 | MSINPELAENTLFLKRHESILKCLNYLIENRQEVMDILTQFSSYRA ANAEIDSTILTLQGALQEVQTYQPSWQRSMAVFMPSNVILYSYAL YLLIPSLYVENIDFRPSSHVNEYVNMLHEKLQAVHGLPIYIRKVS QRVFMENSVMPADIVVFTGSYVNAEKIKKQIRKDQLYIFYGQGIN PFVIGPDADLELAVTDVIRMRLFNSGQDCLGPDIILVHQEVSEHF KDLLIQRLDQLVFGANDDPNANYSPIFYKDALNSVSEYFNTNDK FIIYGGGIDFRTKKMEPTVVYSELDQNLEIIEYFSPVFNVVSYQDD EQLIKRISSSYFSERAMGCSLYGSEHLTDVLRKKHTLTINQTLEDV DQGNKPFGGYGTMSNYIFYDFKLISKPILISEIVAEYLSEKRSLV* |
| ALDH-027 | A0A2E8CRC4 | MIAPHIPILRQGQVYKSLDTQTVNKLGSDEPAAEISFACADMIKY DIQNMGSAREALKKFSGEELVEITKKAGELFLNGDLPIGTEGELQ SPEDYVFSLAATSGLPHSLIRFNMRRLGGLFGQIDDIFKGLSRGLE WSVLDRGYGEQGGAPVSFSPTTNELAVILPSNSPAVNALWIPAIA MKVPVLLKPGREEPWTPWRIIQAFIKAGCPPEAFSLYPTQHDGSG AIIRRAGRVMLFGDDSTVKQYENDERVEVHGTGYSKFVIGEDEIE NWESYIDSMVESVSANSGRSCICTSTIVVPKYGDEIAHALSKKLC EIKPLPQDDPEAKLSGFANPKFAEWIDEAVEEGLQSDGARDITAD YRDGARFVERDGMNYLQPTIVRVDSFDEDLALREFLFPFASVVE CPQSDVVEKTGYSLVMTAVTKDVEWINELFDHPEIERLNIGNVPT NRISWNQPHEGNLFDFLYTRRSFAFAESA* |
| ALDH-028 | A0A2V7BEQ0 | MEPLGVVVVTPPWNFPLSIPAGGVLAALAAGNAVVLKPAPEAVL VGWHLANCLWDAGIPREILQFLPCPDDEIGRGLVTDSRVGGVILT GSAETARLFLGWRPDLPLFAETSGKNAIIITALADRDQAIRDLVRS AFGHNGQKCSAASLAICEAEVYDDADFRRQLRDGAQSLAVGTA WEPTSRITPLTQAPGAALRRALTVLDEGEEWLLEPRPATDNPQLW SPGIKLGVRAGSFFHRTECFGPVLGLMRAENLDHAIELANAQPF GLTSGIQTLDDREIARWVDRIEAGNLYVNRPITGAIVGRQPFGGW KASSVGPGAKAGGPNYVLQLARWRQVARPAVDQEPLSESLATV LDRCLAGLTDADARSLLEASAASYARAWREHFSREHDPSAIRGE LNAFRYRPCRHVIARGMTARPEAAVALCQIILAAHVAGTRLTVSL SPESEPWVGLAECAGVELVVEAEAGFVDRLAHPLYREHGWIERL RAWEPISTAARAAANGTGVTVIDAPVLANGRLELRWYLREQTVS RVLHRYGSVTEPDA* |
| ALDH-029 | A0A1T1H988 | MTASTSSSNTGQCFINGVWQAGEGSEFTSLNPATGEVIWQGKEA SAAQVELAVNAAREASVEWAMMPFADREAIARRFAELLGDNKE EMATIIATETGKPVWETRTEVGAMVGKIAISVNAYNERTGSRVSD VAGARAVLRHKPHGVVAVFGPYNFPGHLPNGHIVPSLLAGNTVL LKPSELTPHVAEFMVQLWEKAGLPAGVLNLLQGQKDTGIALAG HDRIDGLFFTGSSRTGHILHEQFAGHPGKILALEMGGNNPLIIDEV ADMKAAVHETIQSAYITSGQRCTCARRLFVPVGEWGDQFIAQLQ |

| | | |
|---|---|---|
| | | EAVSRIKVGGQFDEDAPFMGSLISEAAADGIAAAQDNLIGLGASP LVKLEKLKPGTGFLSPGLIDVTALVEADKLPDDEYFGPLLQVIRFS DFDEAIRQANNTQYGLSAGLFSDSEARFNYFYDRIRAGIVNWNK QLTGAASSAPFGGVGASGNHRASAYYAADYCSYPVAGMEADHL VLPENLSPGLTIK* |
| ALDH-030 | A0A382Q6S7 | AVHINAFNFPIWGMLEKIAVNLMAGVPAIVKPATLTCYLTELMVR EIIATQILPEGSLQLICGSANGILDHVCCEDVVTFTGSASTGKMLK AHSKLIDEAVPFNMEADSLNASIIGEDAIPGTEEFDLYIKEIQKEM TVKAGQKCTAIRRIIVPEKLVEDVQKALCDRLSKTIIGDPAIEGVR MGSLAGNSQVVEVSEKVNELAESQDIIYGDLENFDVVGANKNK GAFIPPILFLNDNPFEKTDCHIIEAFGPVSTILPYKNLDEAIELARM GKGSLVCSIVTSDDNIAREFTVNAASMHGRILVLNKDCAKESTG HGSPMPLLTHGGPGRAGGG* |
| ALDH-031 | M4Z1V7 | MHTLSASANEVTFIVQRARIAQKCFEGAQQREIDLAVAAAGWRC YRDETAQELSSLAIEETELGNAVDSYQRLRKRILGTLRDLADAVT VGLVHEDVARGVRKFAKPVGVIAGITPATAPAAAAIVNSLSSLKT RNAIIFCPNPRAHRTVGRVVELVRDALREVGAPVDLVQCVGVPT RSISEELMATADLTIASGGASTVRRAYRSGKPALGAGVGNAVVV VDDTADLDAAASMIIAGKSFDYGTSCSSESCILVDESVSDQLIGK LVENGAYMCSGPEMASLRRTAWPDGELAREIVGKSAKQIADRA CIEVPQKTRVLLTLPETTHAEPLGGEKLSPILALWKFAQFDHAVG LVQRLVAASGAGHSCAIHTNVRERAEILARTINVSRVLVNQSTGM GNSGSFDNGLPFSVTLSCGTWGGGSTTDNVNWRHFLNYTWLSE TIPKNEPRPEDLFSEYWSVYHPTASYVGQLGGM* |
| ALDH-032 | A0A420XXS9 | MAQGISIKSPVGDKLDWDCQELKAKFPKTKDDIKEPIPSPESCLL LIDGKVVKSEKTATEVESPVITQDDNKHVIIGKFEMASEAQAMQ ALEAAQRAYGYGREEWSKMKLEERCKHMEGFANDLEKKTDEI AQLLMWEIGKTSKDAKSEVTRTVAYIQTAIKEAKALSESESKWE EEKGILCQIRHLPVGVVLASSPFNYPLNEAYTTFIPALLMGNSVIV RAPRNGATPHFPTLELFAKHFPAGTIQFLTGSGREVMGSLIGTGKI DAVAFIGTASSAAGLFKGAPNPQKLRVMYEGEAKDSAIILPDADL DVAVESCASGMTSFNGQRCTAIKMIWVHESKAEEFLKKLGEKID SMQLGMPWEEGVKITPLCEDTKPGYIKELIEDALKKGAHIVNNG GKCYATLCTLSILAPANKDMKIWSEEQFGPVTPVAFYTDLQEPID YVAKSEYGLQASVYGYDEDDIAKVVDALSYHVGRLNINAPDQR GPDVFPFTGRGNSALGILNAPEGLKFFSVPTMVATKTDDERNVK VLKAVNAGGKSQVVFGAK* |
| ALDH-033 | A0A1F6LNY1 | MASEQEIHQVVGKVMEKVRQRYTVGTGGKIFSSAPGRATSPSM DFRPRKAVGGDSTFDDPDAAAKAARQAQRELMALGLEKRFEIV AAMRAAALENARRLGELAVSETKFGKMPDKMQKVELAARKTP GPEIIHPVTFTGDHGLTLVEKAPYGVMVSITPTTNPPSTVVNNSIG MVSAGNAVIINPHPNAKEVSCEAARILQLAIESAGGPKHLVCAIG NPSQESGTKLMNHPGVDATLVTGGREIVRVAMSSGKKVFAAGPG NPPVVVDETCVLPKAAKDTVFGAQFDNCALCTGEKEIFCVASIC DDFKAEMGKNGSVEIKGRDAERLTKLIIVEDSGSGERHPRVNKE LIAKDAVKILAQAGIRAPAGITHAFLEVDWDHPLVMAEQLMPITP |

| | | |
|---|---|---|
| | | IVRCKDYDQAKEWAILAEHRFRHTFTMHSTNIARLSDMARASN ANIFVKNGPSLAGLGFGGEGYTTLSIAGWTGEGFTTAATFTRERR CTLVDYFRIV* |
| ALDH-034 | A0A7H4GQ81 | MNFEPTGIDPLLDEALSAIATRSAFSGFKESGSSRIHGTEKPAAGK AAYEACLGQAFRLDDSEADFIAAEEVSPFTGEALGIRYAGPDPDR QFAAARAAMKGWASASVEQRIKLCVEMLRQVDEHGFEIAHAIM HTAGQSFPMAYAGSGANALDRGLEAVAWSWLMMNALPFQASW QKQVGAGDPVRLAKRWRLMPRGPAVVFCCASFPTWNGYPAMM ANLACGNPVLVKPHPTAVLPMAIAVRIIRKVLAEAGFDPGLIQLV VDSSSKPLGKVLVEHDQAAIIDFTGSARFGSWVEQHAGNRPCYT ETSGINTVILDGAEDIDAALDSVVNTLCLFSAQMCTSPQTFYVPA SGMSDGNGRCLSADEVEAALAGRIDALAANPKRAASVMGCIQS PATLSLIDEWTGLAESGRYRLLHRSDHYAHPAGPDARTATPLVLG LDLEDGTAPQEEVFGPIAFVIRTADRDQALQHAADSARRHGAITA YLYSTDEDYIESATDAFAWAGAALTCNVTGNMPLNFSAAYSDLH VSGLNPAGNATLAEPSFITGRFRYAQVRRPLSGDGDE* |
| ALDH-036 | A0A3D2UMU3 | MSFPEIKMPELEEVKKHFKLFDYIDGERIAPTVDMKAYVHNPNT GEKIAKQMATSNENIERALQVAQRLHESGEWANTSPEKRAEILD NAANHLMGLVPMIAAVESYNTGIVFSTTSFVNAIVWLAFKAAAG VLKSGYAVTKVPGPNGDVEVLRKPLGVAVCIVPWNSPAALAAH KIANALAAGCPVILKPTEWAPYSCLLLADGLEAAGLPKGLFQIV NGGAEVGNKLVADKRVRAVSFTGGLIGGRAVAEACAKDFKSLQL ELGGNNAMVVLQDADVDKVADGIVTGMTTLNGQWCRALGRL VVHESIQEQVVKAAIEKFKQIKVGDAMSTESQLGPLANEVHKGR IVDSINALKQKGGQTHQSAPLPDSAGFYLSPTLITGVAPEETLEEI FGPVATVHTFKTDDEAIKLANQTPYGLGGYVYSKDESHAMEIAR KMNTGGVKVNGVSLLELNVDAPRPAWGLSGFGEEGTLETFDVF CGKSVVGIAGK* |
| ALDH-037 | P05091 | MLRAAARFGPRLGRRLLSAAATQAVPAPNQQPEVFCNQIFINNE WHDAVSRKTFPTVNPSTGEVICQVAEGDKEDVDKAVKAARAAF QLGSPWRRMDASHRGRLLNRLADLIERDRTYLAALETLDNGKP YVISYLVDLDMVLKCLRYYAGWADKYHGKTIPIDGDFFSYTRHE PVGVCGQIIPWNFPLLMQAWKLGPALATGNVVVMKVAEQTPLT ALYVANLIKEAGFPPGVVNIVPGFGPTAGAAIASHEDVDKVAFTG STEIGRVIQVAAGSSNLKRVTLELGGKSPNIIMSDADMDWAVEQA HFALFFNQGQCCCAGSRTFVQEDIYDEFVERSVARAKSRVVGNP FDSKTEQGPQVDETQFKKILGYINTGKQEGAKLLCGGGIAADRG YFIQPTVFGDVQDGMTIAKEEIFGPVMQILKFKTIEEVVGRANNS TYGLAAAVFTKDLDKANYLSQALQAGTVWVNCYDVFGAQSPF GGYKMSGSGRELGEYGLQAYTEVKTVTVKVPQKNS* |
| ALDH-038 | P51647 | MSSPAQPAVPAPLANLKIQHTKIFINNEWHDSVSGKKFPVLNPATE EVICHVEEGDKADVDKAVKAARQAFQIGSPWRTMDASERGRLL NKLADLMERDRLLLATIEAINGGKVFANAYLSDLGGSIKALKYC AGWADKIHGQTIPSDGDIFTFTRREPIGVCGQIIPWNFPLLMFIWK IGPALSCGNTVVVKPAEQTPLTALHMASLIKEAGFPPGVVNIVPG YGPTAGAAISSHMDVDKVAFTGSTQVGKLIKEAAGKSNLKRVTL |

| | | |
|---|---|---|
| | | ELGGKSPCIVFADADLDIAVEFAHHGVFYHQGQCCVAASRIFVEESVYDEFVRKSVERAKKYVLGNPLTQGINQGPQIDKEQHDKILDLIESGKKEGAKLECGGGRWGNKGFFVQPTVFSNVTDEMRIAKEEIFGPVQQIMKFKSIDDVIKRANNTTYGLAAGVFTKDLDRAITVSSALQAGVVWVNCYMILSAQCPFGGFKMSGNGRELGEHGLYEYTELKTVAMKISQKNS* |
| ALDH-039 | P77674 | MQHKLLINGELVSGEGEKQPVYNPATGDVLLEIAEASAEQVDAAVRAADAAFAEWGQTTPKVRAECLLKLADVIEENGQVFAELESRNCGKPLHSAFNDEIPAIVDVFRFFAGAARCLNGLAAGEYLEGHTSMIRRDPLGVVASIAPWNYPLMMAAWKLAPALAAGNCVVLKPSEITPLTALKLAELAKDIFPAGVINILFGRGKTVGDPLTGHPKVRMVSLTGSIATGEHIISHTASSIKRTHMELGGKAPVIVFDDADIEAVVEGVRTFGYYNAGQDCTAACRIYAQKGIYDTLVEKLGAAVATLKSGAPDDESTELGPLSSLAHLERVGKAVEEAKATGHIKVITGGEKRKGNGYYYAPTLLAGALQDDAIVQKEVFGPVVSVTPFDNEEQVVNWANDSQYGLASSVWTKDVGRAHRVSARLQYGCTWVNTHFMLVSEMPHGGQKLSGYGKDMSLYGLEDYTVVRHVMVKH* |
| ALDH-042 | A4IT08 | MIYAQPGQPGALVTFKKRYENFIGGKWVPPVDGEYFENITPITGKPYCEVPRSKAADIELALDAAHAAKDEWGRTSPAKRARLLNKIADRMEENLELLAVAETWENGKPIRETLAADIPLAIDHFRYFASCIRAQEGTISEIDHDTVAYHFKEPLGVVGQIIPWNFPILMAAWKLAPALAAGNCVVLKPAEQTPTSILVLIELIEDLLPPGVVNIVNGFGLEAGKPLASNPRVAKVAFTGETTTGRLIMQYASQNIVPVTLELGGKSPNIFFADVMDKDDEFFDKALEGFTMFALNQGEVCTCPSRALIHESIYDAFMERALERVKQIKQGNPLDTETMIGAQASSEQLEKILSYIDIGKQEGAELLIGGERNMLEGELAGGYYVKPTIFKGHNKMRIFQEEIFGPVLAVTTFKDEDEALAIANETLYGLGAGVWTRNINTAYRFGRGIQAGRVWTNCYHVYPAHAAFGGYKMSGIGRETHKMMLDHYQQTKNMLVSYSPKKLGLF* |
| ALDH-045 | P51977 | MSSSAMPDVPAPLTNLQFKYTKIFINNEWHSSVSGKKFPVFNPATEEKLCEVEEGDKEDVDKAVKAARQAFQIGSPWRTMDASERGRLLNKLADLIERDRLLLATMEAMNGGKLFSNAYLMDLGGCIKTLRYCAGWADKIQGRTIPMDGNFFTYTRSEPVGVCGQIIPWNFPLLMFLWKIGPALSCGNTVVVKPAEQTPLTALHMGSLIKEAGFPPGVVNIVPGYGPTAGAAISSHMDVDKVAFTGSTEVGKLIKEAAGKSNLKRVSLELGGKSPCIVFADADLDNAVEFAHQGVFYHQGQCCIAASRLFVEESIYDEFVRRSVERAKKYVLGNPLTPGVSQGPQIDKEQYEKILDLIESGKKEGAKLECGGGPWGNKGYFIQPTVFSDVTDDMRIAKEEIFGPVQQIMKFKSLDDVIKRANNTFYGLSAGIFTNDIDKAITVSSALQSGTVWVNCYSVVSAQCPFGGFKMSGNGRELGEYGFHEYTEVKTVTIKISQKNS* |
| ALDH-046 | UPI00019FFBE0 | MLKTNIELKPKVEAFLNEEIKMFINGEFVSAIGGKTFETYNPATEDVLAVVCEAQEEDIDAAVKAARSAFESGPWAEMTTAERAHLIYKLADLIEEHREELAQLEALDNGKPYQVALDDDISATVENYRYYAGWTTKIIGQTIPISKDYLNYTRHEPVGVVGQIIPWNFPLVMSSWKMGAALATGCTIVLKPAEQTPLSLLYAAKLFKEAGFPNGVVNFVPGF |

| | | |
|---|---|---|
| | | GPEAGAAIVNHHDIDKVAFTGSTVTGKYIMRQSAEMIKHVTLEL GGKSPNIILEDADLEEAINGAFQGIMYNHGQNCSAGSRVFVHRK HYETVVDALVKMANNVKLGAGMEKETEMGPLVSKKQQERVLN YIEQGKKEGATVAAGGERALEKGYFVKPTVFTDVTDDMTIVKEE IFGPVVVVLPFDSTEEVIERANNSSYGLAAGVWTQNIKTGHQVA NKLKAGTVWINDYNLENAAAPFGGYKQSGIGRELGSYALDNYT EVKSVWVNIK* |
| ALDH-047 | Q9LRI6 | MAARRAASSLLSRGLIARPSAASSTGDSAILGAGSARGFLPGSLH RFSAAPAAAATAAATEEPIQPPVDVKYTKLLINGNFVDAASGKTF ATVDPRTGDVIARVAEGDAEDVNRAVAAARRAFDEGPWPRMTA YERCRVLLRFADLIEQHADEIAALETWDGGKTLEQTTGTEVPMV ARYMRYYGGWADKIHGLVVPADGPHHVQVLHEPIGVAGQIIPW NFPLLMFAWKVGPALACGNAVVLKTAEQTPLSALFVASLLHEAG LPDGVLNVVSGFGPTAGAALSSHMGVDKLAFTGSTGTGKIVLEL AARSNLKPVTLELGGKSPFIVMDDADVDQAVELAHRALFFNQG QCCCAGSRTFVHERVYDEFVEKARARALQRVVGDPFRTGVEQG PQIDGEQFKKILQYVKSGVDSGATLVAGGDRAGSRGFYIQPTVFA DVEDEMKIAQEEIFGPVQSILKFSTVEEVVRRANATPYGLAAGVF TQRLDAANTLARALRVGTVWVNTYDVFDAAVPFGGYKMSGVG REKGVYSLRNYLQTKAVVTPIKDAAWL* |
| ALDH-048 | C9DIJ2 | MLRTATRTTFKSAQPSFMAAAAALRYYSHYPLSKSITLPNGKVY EQPTGLFINGEFVASQQHKTFEVINPSNEDEICHVYEARTEDVDA AVDAAYNAFHSEWSKMDPSIRGEHLMKLAELMEKNKDTLAAIE SMDNGKALFMAEIDVKLVINYLKYCAGWADKLFGKVVDTGSN YFNYIKREPIGVCGQIIPWNFPLLMWSWKVGPALAAGNTIVLKT AESTPLSALYAAKLAKEAGIPDGVINIVSGFGKITGEAISTHPKIKK LAFTGSTATGKHIMKAAAESNLKKVTLELGGKSPNIVFNDADIK KAVNNLILGIFFNSGEVCCAGSRVFIQEGVYDQVLEEFKIAAEAL KVGNPFEEGVFQGAQTSQQQLTKILGYVESGKDEGATLVTGGER LGDKGYFVKPTIFADVKPNMKIYSEEIFGPFAVVTKFKTADEAIA MANDSEYGLAAGIHTTSLDTATYVANNLEAGTVWINTYNDFHH NMPFGGFKQSGIGREMGEAAFENYTQWKTVRIAINDGPQ* |
| ALDH-049 | Q65NX0 | MSVAAESKTYFNFINGRWVKAESGGMEQSLNPADTRDIVGLVQ KSSIEDVDRAVEAAKQAKKAWRKLSGAERGQFLYKAADIMEQR LDEIAECATREMGKTLPEAKGETARGIAILRYYAGEGLRKTGDVI PSTDSSAFMYTDRVPLGVVGVISPWNFPVAIPIWKMAPALIYGNT VVIKPATETAVTCLKVISCFEEAGIPSGVVNAVTGPGSSAGQRLAE HPDVNGITFTGSNQTGKIIGRTAFERGAKYQLEMGGKNPVIVAD DADLDIAVEAVISGAFRSTGQKCTATSRVIVLNGVYDRFKEKLLQ QTKEITIGDSLKEDVWMGPIANKQQLDNCLSYIAKGKQEGADLI FGGERLADGKYENGYYIRPAIFDNVTSGMTIAQEEIFGPVIALIKA DTLEEALETANDVKFGLSASIFTQNIRRMLSFTDEIEAGLIRVNAE SAGVELQAPFGGVKQSSSHSREQGEAAKEFFTAVKTVFVKP* |
| ALDH-050 | UPI00005BF137 (*Bt*ALDH3) | MSAISEVVQRARAAFNSGRTRPLQFRVQQLEGLRRLIREREKDLV GALAADLHKNEWTAYYEEIVYVLEEIDYMIRKLPEWAADEPVE KTPHTQQDEAYIHSEPLGVVLIIGSWNYPFNLTIQPMVGAIAAGN |

| | | |
|---|---|---|
| | | AVVLKPSELSENTASLLATILPQYLDQDLYPVINGGVAETTEVLKE<br>RFDHILFTGSTGVGRVVMMAAAKHLTPVTLELGGKNPCYVDKD<br>CDLDIACRRIAWGKFMNSGQTCVAPDYILCDPSIQSQVVEKLKK<br>SLKEFYGEDAKKSRDYGRIINSRHFQRVMGLLEGQKVTYGGTG<br>DATTRYIAPTILTDVDPESPVMQEEVFGPVLPIMCVRSLEEAIQFIT<br>QREKPLALYVFSPNDKVIKKMIAETSSGGVTANDVVVHISVHSLP<br>YGGVGDSGMGSYHGRKSFETFSHRRSCLVRPLLNEETLKARYPP<br>SPAKMPRH* |
| ALDH-051 | P20000 | MLRAVALAAARLGPRQGRRLLSAATQAVPTPNQQPEVLYNQIFIN<br>NEWHDAVSKKTFPTVNPSTGDVICHVAEGDKADVDRAVKAARA<br>AFQLGSPWRRMDASERGRLLNRLADLIERDRTYLAALETLDNG<br>KPYIISYLVDLDMVLKCLRYYAGWADKYHGKTIPIDGDYFSYTR<br>HEPVGVCGQIIPWNFPLLMQAWKLGPALATGNVVVMKVAEQTP<br>LTALYVANLIKEAGFPPGVVNVIPGFGPTAGAAIASHEDVDKVAF<br>TGSTEVGHLIQVAAGKSNLKRVTLELGGKSPNIIMSDADMDWAV<br>EQAHFALFFNQGQCCCAGSRTFVQEDIYAEFVERSVARAKSRVV<br>GNPFDSRTEQGPQVDETQFKKVLGYIKSGKEEGAKLLCGGGAA<br>ADRGYFIQPTVFGDVQDGMTIAKEEIFGPVMQILKFKSMEEVVG<br>RANNSKYGLAAAVFTKDLDKANYLSQALQAGTVWVNCYDVFG<br>AQSPFGGYKLSGSGRELGEYGLQAYTEVKTVTVRVPQKNS* |
| ALDH-052 | P23883 | MNFHHLAYWQDKALSLAIENRLFINGEYTAAAENETFETVDPVT<br>QAPLAKIARGKSVDIDRAMSAARGVFERGDWSLSSPAKRKAVL<br>NKLADLMEAHAEELALLETLDTGKPIRHSLRDDIPGAARAIRWY<br>AEAIDKVYGEVATTSSHELAMIVREPVGVIAAIVPWNFPLLLTCW<br>KLGPALAAGNSVILKPSEKSPLSAIRLAGLAKEAGLPDGVLNVVT<br>GFGHEAGQALSRHNDIDAIAFTGSTRTGKQLLKDAGDSNMKRV<br>WLEAGGKSANIVFADCPDLQQAASATAAGIFYNQGQVCIAGTRL<br>LLEESIADEFLALLKQQAQNWQPGHPLDPATTMGTLIDCAHADS<br>VHSFIREGESKGQLLLDGRNAGLAAAIGPTIFVDVDPNASLSREEI<br>FGPVLVVTRFTSEEQALQLANDSQYGLGAAVWTRDLSRAHRMS<br>RRLKAGSVFVNNYNDGDMTVPFGGYKQSGNGRDKSLHALEKF<br>TELKTIWISLEA* |
| ALDH-053 | A0A6H1TS81 | MLTATPVREIIERQRQFFATGRTKSVDFRIEQLKKLKQAILDHEAE<br>IIAAVQADLRKPHLEAYLTEIGSVSKIDYALKQIKSWVKPQKVATG<br>IEQFPASARVYSEPLGVVLIISPWNYPFNLAIEPLIGAIAAGNCAIV<br>KPSEVSANTSRVIAKLFGAVFDPGYISVVEGDAEVSQGLLAEKFD<br>HIFFTGGTAIGQRVMEAAAKQLTPVTLELGGKSPCIVDPQINLETA<br>ATRVTWGKFLNAGQTCIAPDYLLVDRRIQADFVAEIQKKLHQFF<br>GPSPQESPDFGRIVSDKHFQRLASLLQDAQIVTGGELDPGDRYIA<br>PTLVENVALDAPLMQEEIFGPILPIIPYDRFDEAIAIVNQRPKPLAL<br>YLFSNDKEKQARIVRETSSGGVCLNDTIMHVGVAELPFGGVGPS<br>GIGAYHGKASFDTFSHRKSVLKKSFWLDLDLRYPPYAGKLKLVK<br>KFLGQ* |
| ALDH-054 | A0A7L1RJ35<br>+A0A7L1RY59† | MERMQQIVGRARAAFNSGRSRPLEFRIQQLKALERMVQEKEKEI<br>LAALKADLNKCGHNAYSHEILGVLGELAQTMEKLPSWAAPQPV<br>KKNLLTMRDEAYINYEPLGVVLVIGAWNYPFVLVMQPLIGAIAA<br>GNAVVVKPSEVSENTAQLVAELLPQYLDKELYPVVTGGVPETTE<br>LLTQRFDHILYTGNTAVGKIVMAAAAKHLTPVTLELGGKSPCYID |

| | | |
|---|---|---|
| | | KDCDLAVACRRITWGKYMNCGQTCIAPDYILCDPSIQGKVVENIKATLKEFYGEDVKSSPDYGRIVSQRHFKRVMSLLEGQKIAHGGETDEASCFIAPTILTDVSPESKVMEEEIFGPVLPIVTVRSVEEAIEFINRREKPLALYVFSNNKQLIKRVISETSSGGVTGNDVIMHFFLSTLPFGGVGHSGMGAYHGKHSFETFSHRRACLIKDLKMESTNKMRYPPGSQKKV* |
| ALDH-135 | A0A662CPS5 | MKVPWASLDERLEVLADMGRRIEARREDFLEALALDIGQAVKVTSEEIRLTMEHLSTMEEEASLLEGREPYGLVGAIFPYDGPTVMFARWGGAALLGGNRLRFSFSSLTPRVAQLMEEVCQPWKDVVEVVIGKDNREFGWDCVEDPDVRVFFVSGSREVGRVYAQAIDEFDKVIFAGPGGMPPVLVFSGASVERAAVFAARRAFLNGGQYCTTIKRALVHKDLLDPFVEALLEEVDKIKVGDPMDPQVDYGPIKAERTRVLFERGLEWVKGTLIRGGPPEGEWIFPTVVLAREIPDIEVFGPFLAVKAMGSDRAMVEEAVRTGYPLIAYAFGPPPLGSKGRLEALYGRVYWDPEFLYLSPRDPFGGRRDSGWVLEKRGAMIIRKGGPIVYVEELTRPVSGSSG* |
| ALDH-136 | A0A1J5KC07 | MTEKLKCLNARTGEVMEELAITPVSEIKNIVAKSHTAQKKWALLSIDERSDYIRKAYETITEDLEGFANLIHEEMGKTMPEALGEIKVYTSGLENMIKEVKEALTPEVNSAGELETTTYFDSLGVCASITPWNFPMGMPHTLMMPSLMAGNTIVFKPSEEVTLIGLAYAKYLNKFLPEDVLQVVVGAGEQGKALVEADVQLVTFTGSQRTGKNILETGAKDLKRVILELGGKDALIVMDDVDIDSASKFATINSFRNCGQVCVSTEKILVTDKNHDIFVEKLVERAKRLEVSSLINKTQKNHVLAQIDDAMKKGAKMVLGDPSKDEKNFLSPVVLTNVTNDMDIMIDETFGPVACITKVNDLDHAVELANEGEYALGSVIFGNDKEQAKNVARRLKAGMIGINKSCGGVKGSPWVGAGQSGYNFHGSSAGHRMFTQVRIVTA* |
| ALDH-137 | A0A3L7TS19 | MRAINPATGESFGDEFARSTRDELAQMAEAALEVVDVLADAPSAQLAGFLDDFAMRLESDRDEIAAIAHAETALPLTPRLREIEFNRTTGQLRMAANGLRDESWREVAVDANNNLRSTLVPLGGAVLCLGPCNFPLAYNGVSGGDFCSAIMARNPVIAKAHPSHPNTTARMFAHAVAARDAASLPPASVQMFFDCANEDGEALLAHRGVAALGFTGSRAAGVRLKSVCDALGKPASLEMASINPVFVASNALTARHDAIADAWTASLLMAGGQQCTKPGVIFVSGIDAAARFIARAEHNARAVAPAVLLSESIRTYLSDSISAWKNVGASIRCGGNASSPGIRWEPTLLSVDAAFAALHRDLLHHEAFGPLGVIVICDSDAQLVEFARALDGQLAGTIVSDASDANTRHALQRALRFKVGRMLDEAMPTGVVVSNAMVHGGPFPATGDARFTAVGLPASAKRFSKTLCWDRCRA* |
| ALDH-138 | A0A497RVJ2 | MKEFPNFLHYKGKINRIYPEDRESEGAIVIKDYLTEEEIGYFPDLKKVSKAIGGARNLHKYWNRLEKKDRVEILEQAGEELEKKNELDKLISRAGGFPIRFVREARENFADYLKQSEEFLQENPGKGPVVAATSVTTPEVQPYVMIESLLGNCSVTIKGNSSEPFSAYLLSEISEETDLPIQFITYQTKGKESFATEFYKMCEEEGGHFILMGDPITPKRIAYYEILDKVDISTLPMPKNMVAFTSHGGCMIVDESGSIEEAVEGALYSFRFPKACKVPTCIFVHKDRVEEFADELVGKVKKQKVGDVLDEDTEIAEVSEKYWEELVEPFLRVAKSEGETLVGGDINQPSVIKGNFYLSLWMEPHFPIYCIEEFEDVREKIEKINKAGKNLGGKILDLSVYSDDQFFEELEKLRREGSLRVYTLHKNSPTTSFDPKTAHEGIILREYLAEPNFVQK* |
| ALDH-139 | A0A3D5C0Z5 | MASNRGLVYLGPHNVEVQTTDYPELSLGTRSCQHGVILKVVATNICGSDQHMVRGRTTAPSGLVLGHEITGQVVEAGQDVEFISVGDIVSVPFNIACGRCRNCKEGKTGICLHVNPARPGAAYGYVDMGGWVGGQAEYVMVPYADFNLLKFPDADQALEKIRDLTLLSDIFPTGFHGAVSAGVGPGSIVYIAGGGPVGLASAAGSQLLGAALVIVGDMIPERLAQARSFGCETVDLTQDATLAEQIDQIVGEPEVDSAVDCVGFEARGHGSESNVERPATVLNSLMEVTRAGGSIGIPGLYVTGDPGGV |

| | | |
|---|---|---|
| | | DENAKVGNLGIRIGLGWAKSHDFTTGQCPVMRYHRPLMQAILH<br>EKTRPAEAVNVTVISLDEAPKGYKDFDKGAAKKYVIDPHGMVA<br>V* |
| ALDH-140 | A0A2A5G690 | MTQDIETVESQPQQPVSSGNPFAEIVRKQRDFFNSGASKSAEFRI<br>KQLKNLKEAFHKHHENLEKALYEDLRKSKTEAFATEIGIMIAEIE<br>HNMKNIRKWMQPKKVKTPLFFMPGKSRIHYEPFGVSLIISPWNY<br>PVKNLFGPVLGAMTAGNCSVLKPSEVSPYTSAVAKKMVDEFFDP<br>EFMTVVEGGVPETSELLKLKWDYIFFTGGTEIGRIIYQAAAKQLA<br>PCTLELGGKSPTIVDKDINLDVTAKRLCWGKFVNAGQTCVAPDY<br>LLVHKDIKQKLIEKLKETINEFYGENPAESPDLGRIISDRHYNRIK<br>NLIDGDVIFGGQCDESQRYIAPTIIDNVSPDAKVMQQEIFGPILPII<br>EYENVDEAIAFINDREKPLALYLFSNNENVRRKIIDNTSSGGVCIN<br>ETIMHMASPEMPFGGVGNSGMGAYNGRFGFDTFSHKKPVMTRS<br>FLFDVKQKYAPFNAKKLNFVKFALKRLI* |
| ALDH-141 | A0A1Q5PDQ4 | MSHTNTIRTSPATAETNLHRIQALVKKQREFFSSGHTLDLAFRKE<br>RLRELQHTIQEHEQDLMDAMYADFHKPEMEAFSTEIGFVELELK<br>LVLKNLQKWAKPKRVKESLLNFPSRSYIHSDPFGVALIIGPWNYP<br>FQLLLNPLIGAMAAGNCAIVKPSELTPTTSAVVARMIRQHFDESYI<br>ATVEGGAPTTQHLLQQRFDYIFFTGSTQVGKIVMKAAAEHLTPV<br>TLELGGKSPAIVAEDADLGLAARRIAWGKFLNAGQTCIAPDYLL<br>AQESIKEELLQLISQCIRDFYGEDPRQSPDFARIVNDRHFNRLSGF<br>MSEGKVRTGGVTDAASRYIAPTLLDRVTWQHPIMQEEIFGPILPV<br>LSFQHLDEAIGMVQQREKPLALYFFTNDERKKEQVLKYTSFGGG<br>CINDTISHIINPNLPFGGVGQSGMGSYHGQSSFELFSQQKSVLHR<br>GTWLDLPLRYPPYGNRLPMLRKFFKWL* |
| ALDH-174 | Q84DC3 (*Pp*MdlD) | MNYLSPAKIDSLFSAQKAYFATRATADVGFRKQSLERLKEAVINN<br>KEALYSALAEDLGKPKDVVDLAEIGAVLHEIDFALAHLDEWVAP<br>VSVPSPDIIAPSECYVVQEPYGVTYIIGPFNYPVNLTLTPLIGAIIGG<br>NTCIIKPSETTPETSAVIEKIIAEAFAPEYVAVIQGGRDENSHLLSLP<br>FDFIFFTGSPNVGKVVMQAAAKHLTPVVLELGGKCPLIVLPDAD<br>LDQTVNQLMFGKFINSGQTCIAPDYLYVHYSVKDALLERLVERV<br>KTELPEINSTGKLVTERQVQRLVSLLEATQGQVLVGSQADVSKR<br>ALSATVVDGVEWNDPLMSEELFGPILPVLEFDSVRTAIDQVNKH<br>HPKPLAVYVFGKDMDVAKGIINQIQSGDAQVNGVMLHAFSPYLP<br>FGGIGASGMGEYHGHFSYLTFTHKKSVRIVP* |
| ALDH-178 | P30838 (*Hs*ALDH3) | MSKISEAVKRARAAFSSGRTRPLQFRIQQLEALQRLIQEQEQELV<br>GALAADLHKNEWNAYYEEVVYVLEEIEYMIQKLPEWAADEPVE<br>KTPQTQQDELYIHSEPLGVVLVIGTWNYPFNLTIQPMVGAIAAGN<br>SVVLKPSELSENMASLLATIIPQYLDKDLYPVINGGVPETTELLKE<br>RFDHILYTGSTGVGKIIMTAAAKHLTPVTLELGGKSPCYVDKNC<br>DLDVACRRIAWGKFMNSGQTCVAPDYILCDPSIQNQIVEKLKKSL<br>KEFYGEDAKKSRDYGRIISARHFQRVMGLIEGQKVAYGGTGDAA<br>TRYIAPTILTDVDPQSPVMQEEIFGPVLPIVCVRSLEEAIQFINQRE<br>KPLALYMFSSNDKVIKKMIAETSSGGVAANDVIVHITLHSLPFGG<br>VGNSGMGSYHGKKSFETFSHRRSCLVRPLMNDEGLKVRYPPSPA<br>KMTQHSRGIK* |

†To clone a complete ALDH-054 from fragment sequences, A0A7L1RJ35 (residues 1-112) was concatenated to A0A7L1RY59 (residues 1-339).

**Table S3:** Colorimetric Screening Data

| Plasmid # | UniProt ID | Substrate Tested | Activity (nmol Formazan $mg^{-1}$ $min^{-1}$) | |
|---|---|---|---|---|
| | | | $NAD^+$ | $NMN^+$ |
| *Bt*ALDH3 (ALDH-050) | UPI00005BF137 | Hexanal | 27.83 | 13.95 |
| *Pb*ALDH (ALDH-010) | L8N0N6 | Hexanal | 657.98 | 13.43 |
| ALDH-039 | P77674 | Butyraldehyde | 119.49 | 10.66 |
| ALDH-013 | A0A2Z4LU87 | Butyraldehyde | 134.67 | 6.89 |
| ALDH-049 | Q65NX0 | Hexanal | 688.35 | 3.19 |
| ALDH-009 | A0A0D6KE96 | Butyraldehyde | 48.65 | 2.19 |
| ALDH-046 | UPI00019FFBE0 | Butyraldehyde | 26.29 | 1.61 |
| ALDH-014 | A0A0S8BPB2 | Hexanal | 58.61 | 1.17 |
| ALDH-137 | A0A3L7TS19 | Butyraldehyde | 20.48 | 0.97 |
| ALDH-048 | C9DIJ2 | Acetaldehyde | 527.99 | 0.88 |
| ALDH-037 | P05091 | Hexanal | 59.11 | 0.72 |
| ALDH-045 | P51977 | Hexanal | 42.73 | 0.69 |
| ALDH-017 | A0A517R638 | Hexanal | 44.27 | 0.5 |
| ALDH-027 | A0A2E8CRC4 | Hexanal | 75.69 | 0.37 |
| ALDH-052 | P23883 | Butyraldehyde | 588.93 | 0.34 |
| ALDH-038 | P51647 | Hexanal | 235.91 | 0.21 |
| ALDH-042 | A4IT08 | Acetaldehyde | 16.46 | 0.1 |
| ALDH-029 | A0A1T1H988 | Hexanal | 300.63 | 0 |
| ALDH-136 | A0A1J5KC07 | Acetaldehyde | 201.78 | 0 |
| ALDH-033 | A0A1F6LNY1 | Acetaldehyde | 80.06 | 0 |
| ALDH-030 | A0A382Q6S7 | Acetaldehyde | 70.31 | 0 |
| ALDH-016 | A0A7Y1V182 | Butyraldehyde | 43.03 | 0 |
| ALDH-023 | A0A1X6ZLY8 | Hexanal | 42.2 | 0 |
| ALDH-138 | A0A497RVJ2 | Hexanal | 40.95 | 0 |
| ALDH-034 | A0A7H4GQ81 | Hexanal | 40.34 | 0 |
| ALDH-047 | Q9LRI6 | Acetaldehyde | 39.8 | 0 |
| ALDH-021 | UPI000A40ADAD | Hexanal | 38.93 | 0 |
| ALDH-020 | A0A2E3KN37 | Acetaldehyde | 27.82 | 0 |
| ALDH-139 | A0A3D5C0Z5 | Acetaldehyde | 20.75 | 0 |
| ALDH-028 | A0A2V7BEQ0 | Acetaldehyde | 13.16 | 0 |
| ALDH-011 | A0A2V7VB71 | Hexanal | 11.23 | 0 |
| ALDH-024 | A0A060QGV9 | Acetaldehyde | 9.07 | 0 |
| ALDH-015 | UPI000E14A9E8 | Hexanal | 8.98 | 0 |
| ALDH-036 | A0A3D2UMU3 | Hexanal | 7.89 | 0 |
| ALDH-051 | P20000 | Acetaldehyde | 7.2 | 0 |
| ALDH-022 | A0A139NAP6 | Hexanal | 4.3 | 0 |
| ALDH-031 | M4Z1V7 | Hexanal | 3.64 | 0 |
| ALDH-135 | A0A662CPS5 | Hexanal | 2.3 | 0 |
| ALDH-025 | UPI0001E31496 | Butyraldehyde | 2.04 | 0 |
| ALDH-032 | A0A420XXS9 | Butyraldehyde | 2.02 | 0 |
| ALDH-012 | UPI000407F0B2 | Acetaldehyde | 1.29 | 0 |
| ALDH-019 | A0A346XWA7 | Butyraldehyde | 0.54 | 0 |

**Table S4:** Data Collection and Refinement Statistics

| | *Bt*ALDH3 ($NAD^+$) | *Bt*ALDH3 ($NMN^+$) |
|---|---|---|
| Data Collection and Refinement Statistics | | |
| Wavelength (Å) | 0.9793 | 0.9793 |
| Resolution range (Å) | 40.98 – 1.46 (1.51 – 1.46) | 92.86 - 1.46 (1.51 - 1.46) |
| Space group | P1 | P 1 $2_1$ 1 |
| Unit cell dimensions (Å, °) | 46.68, 61.53, 87.95; 94.01°, 100.94°, 115.36° | 93.87, 58.98, 158.15; 90°, 98.41° |
| Unique reflections | 142,671 (13,679) | 289,916 (27,983) |
| Completeness (%) | 95.98 (92.15) | 97.81 (94.66) |
| Wilson B-factor ($Å^2$) | 13.59 | 13.42 |
| Reflections used in refinement | 142,324 (13,640) | 289,782 (27,907) |
| Reflections used for R-free | 1,999 (191) | 14,636 (1,469) |
| R-work | 0.1707 (0.3267) | 0.2112 (0.3387) |
| R-free | 0.1967 (0.3611) | 0.2426 (0.3601) |
| No. of non-hydrogen atoms | 7,719 | 15,278 |
| Macromolecules | 6,931 | 13,834 |
| Ligands | 201 | 313 |
| Solvent | 671 | 1,267 |
| Protein residues | 900 | 1,797 |
| R.M.S. deviations | | |
| Bond lengths (Å) | 0.009 | 0.009 |
| Bond angles (°) | 1.03 | 1.13 |
| Ramachandran Plot (%) | | |
| Favored | 98.66 | 98.54 |
| Allowed | 0.89 | 1.01 |
| Outliers | 0.45 | 0.45 |
| Rotamer outliers (%) | 0.13 | 0.07 |
| Clashscore | 3.45 | 4.51 |
| B-factors ($Å^2$) | | |
| Average B-factor | 15.73 | 16.92 |
| Macromolecules | 14.69 | 16.19 |
| Ligands | 14.89 | 25.18 |
| Solvent | 23.81 | 23.79 |

*Values in parentheses are for highest-resolution shell

**Table S5:** Apparent Kinetic Parameters of *Hs*ALDH3 and *Pp*MdlD with $NAD^+$

| | $k_{cat}$ ($s^{-1}$) | $K_M$ (mM) | $k_{cat}/K_M$ ($s^{-1}$ $mM^{-1}$) |
|---|---|---|---|
| *Hs*ALDH3 | 14.11±0.35 | 0.14±0.01 | 103.43±3.56 |
| *Pp*MdlD | 120.01±2.63 | 0.62±0.03 | 193.8±7.93 |

Reactions were carried out in 100 mM potassium phosphate buffer pH 7.4, 5 mM hexanal, 5 mM dithiothreitol, and varying concentrations of $NAD^+$ (0.005 mM-100 mM), and varying concentrations of purified ALDH (0.01-0.5 mg $mL^{-1}$). All reactions were carried out with $n = 3$ biologically independent replicates, except for *Hs*ALDH3 which was carried out with $n = 2$. Data is reported mean ± standard deviation.

### SUPPLEMENTARY METHODS

#### Synthesis and Characterization of 1-benzylnicotinamide bromide ($BNA^+$ $Br^-$)

1-benzyl nicotinamide bromide (1-benzyl-3-carbamoylpyridin-1-ium bromide, $BNA^+$ $Br^-$) was chemically synthesized using the general procedure reported in King et al[1]. (Bromomethyl)benzene was obtained from Sigma. $^1$H NMR (498 MHz, DMSO) δ 9.68 (s, 1H), 9.34 (d, *J* = 6.0 Hz, 1H), 8.99 (d, *J* = 8.1 Hz, 1H), 8.62 (s, 1H), 8.30 (t, *J* = 7.2 Hz, 1H), 8.19 (s, 1H), 7.59 (d, *J* = 6.9 Hz, 2H), 7.45 (d, *J* = 7.1 Hz, 3H), 5.95 (s, 2H).

$^1$H NMR of 1-benzyl nicotinamide bromide ($BNA^+$ $Br^-$)

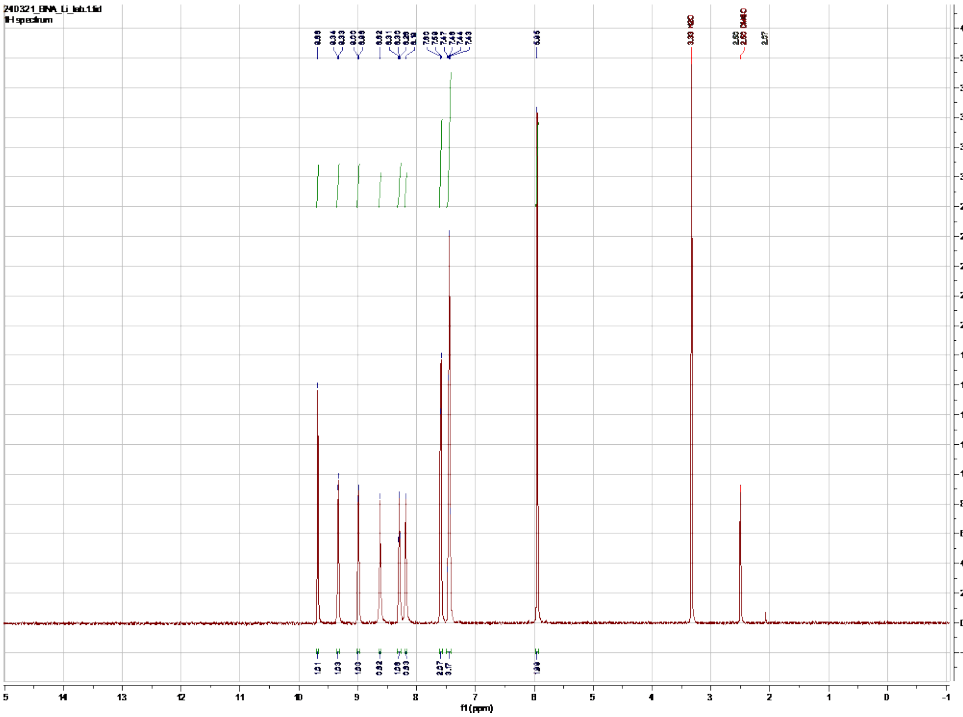


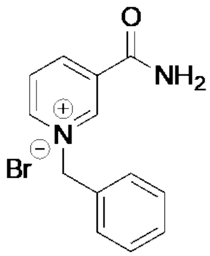

**Synthesis and Characterization of 1-(4-carboxy)benzylnicotinamide bromide (BANA$^+$ Br$^-$)**

1-(4-carboxy)benzylnicotinamide bromide (3-carbamoyl-1-(4-carboxybenzyl)pyridin-1-ium bromide, $^+$BANA Br$^-$) was chemically synthesized using the general procedure reported in Zhou et al[2] with the following starting materials and modifications: nicotinamide (284.54 mg, 2.33 mmole) 4-(bromomethyl)benzoic acid (500 mg, 2.33 mmole). 4-(bromomethyl)benzoic acid was obtained from Sigma. The product was washed with ethyl acetate instead of diethyl ether. The title compound was isolated as a fine powder in 54% yield (1.26 mmole, 426.5 mg). $^1$H NMR (498 MHz, DMSO) δ 9.69 (d, *J* = 1.5 Hz, 1H), 9.35 (dt, *J* = 6.1, 1.3 Hz, 1H), 9.03 (dt, *J* = 8.1, 1.5 Hz, 1H), 8.64 (s, 1H), 8.33 (dd, *J* = 8.1, 6.1 Hz, 1H), 8.21 (s, 1H), 8.04 – 7.98 (m, 2H), 7.71 – 7.66 (m, 2H), 6.06 (s, 2H).

$^1$H NMR of 1-(4-carboxy)benzylnicotinamide bromide (BANA$^+$ Br$^-$)

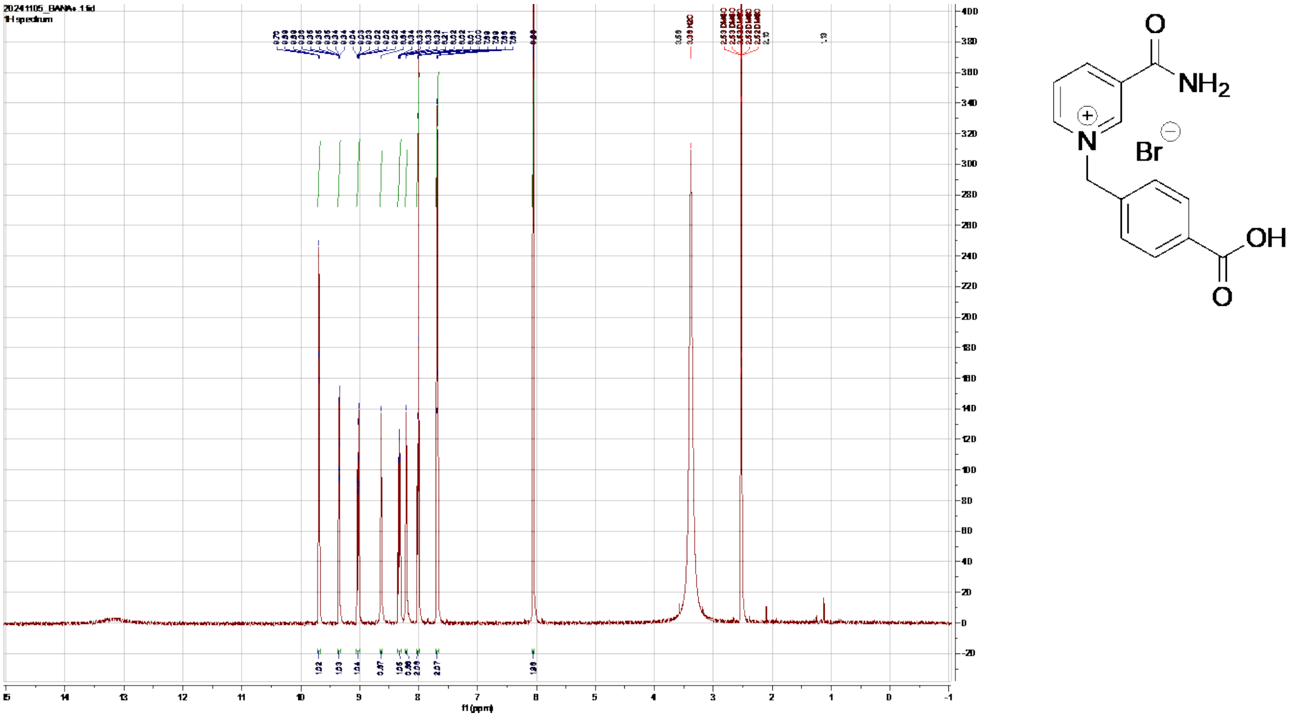

**Synthesis and Characterization of 1-phenylethylnicotinamide bromide (P2NA$^+$ Br$^-$)**

1-phenylethylnicotinamide bromide (3-carbamoyl-1-phenethylpyridin-1-ium bromide, P2NA$^+$ Br$^-$) was chemically synthesized using the general procedure reported in King et al[1] with the following starting materials: nicotinamide (517 mg, 4.23 mmole) 2-phenethyl bromide (2036 mg, 11 mmole). 2-phenetyhl bromide was obtained from Sigma. The reaction mixture was heated to 110°C for 48 hours. The title compound was isolated as a fine powder in 68% yield (2.83 mmole, 869.3 mg). $^1$H NMR (498 MHz, DMSO) δ 9.57 (d, *J* = 1.8 Hz, 1H), 9.18 (dt, *J* = 6.1, 1.3 Hz, 1H), 8.96 (dt, *J* = 8.1, 1.5 Hz, 1H), 8.61 (s, 1H), 8.26 (dd, *J* = 8.1, 6.1 Hz, 1H), 8.18 (s, 1H), 7.37 – 7.24 (m, 5H), 4.95 (t, *J* = 7.7 Hz, 2H), 3.34 (t, *J* = 7.6 Hz, 7H).

$^1$H NMR of 1-phenylethylnicotinamide bromide (P2NA$^+$ Br$^-$)

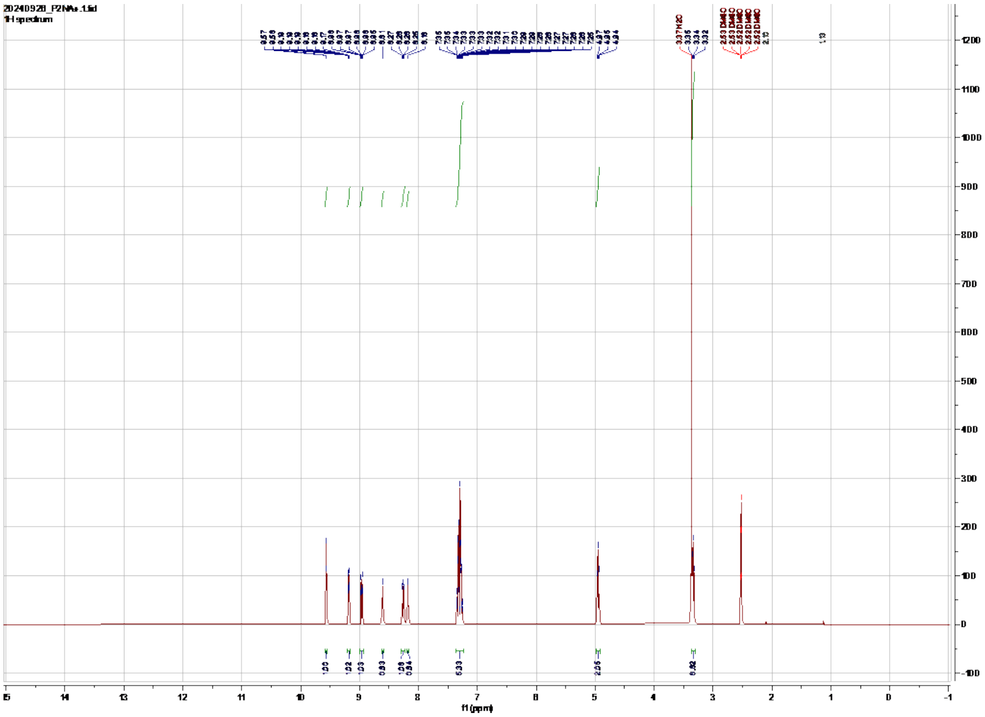


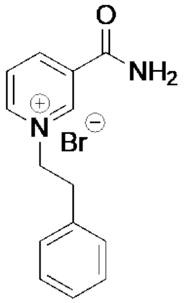

**Synthesis and Characterization of 1-(3-phenyl)propylnicotinamide bromide (P3NA$^+$ Br$^-$)**

1-(3-phenyl)propylnicotinamide bromide (3-carbamoyl-1-(3-phenylpropyl)pyridin-1-ium bromide, P3NA$^+$ Br$^-$) was chemically synthesized using the general procedure reported in King et al[1] with the following starting materials: nicotinamide (441.6 mg, 3.62 mmole) 1-Bromo-3-phenyl propane (720 mg, 3.62 mmole). 1-Bromo-3-phenyl propane was obtained from Sigma. The title compound was isolated as a fine powder in 21% yield (0.76 mmole, 242.7 mg). $^{1}$H NMR (498 MHz, DMSO) δ 9.55 (s, 1H), 9.27 (d, *J* = 6.1 Hz, 1H), 8.95 (d, *J* = 8.1 Hz, 1H), 8.60 (s, 1H), 8.26 (dd, *J* = 8.1, 6.0 Hz, 1H), 8.18 (s, 1H), 7.29 (t, *J* = 7.4 Hz, 2H), 7.27 – 7.22 (m, 2H), 7.20 (t, *J* = 7.2 Hz, 1H), 4.75 (t, *J* = 7.4 Hz, 2H), 2.73 – 2.67 (m, 2H), 2.33 (p, *J* = 7.7 Hz, 2H).

$^{1}$H NMR of 1-(3-phenyl)propylnicotinamide bromide (P3NA$^+$ Br$^-$)

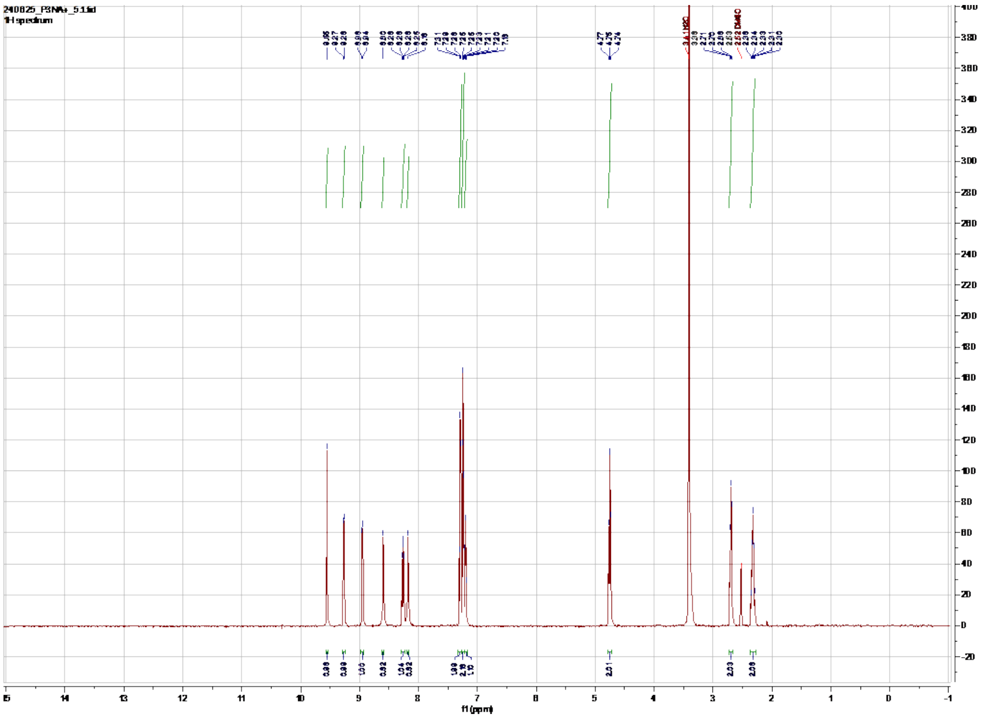

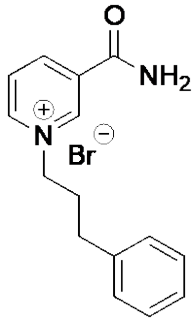

#### Synthesis and Characterization of 1-(3-(4-methoxyphenyl))propylnicotinamide bromide (P3NA-OMe$^+$ Br$^-$)

1-(3-(4-methoxyphenyl))propylnicotinamide bromide (3-carbamoyl-1-(3-(4-methoxyphenyl)propyl)pyridin-1-ium bromide P3NA-OMe$^+$ Br$^-$) was chemically synthesized using the general procedure reported in King et al[1] with the following starting materials: nicotinamide (244.24 mg, 2 mmole) 1-(3-bromopropyl)-4-methoxybenzene (1053 mg, 4.6 mmole). 1-(3-bromopropyl)-4-methoxybenzene was obtained from Ambeed. The reaction mixture was heated to 110$^o$C for 48 hours. The title compound was isolated as a fine powder in 50% yield (1 mmole, 363.7 mg). $^1$H NMR (498 MHz, DMSO) δ 9.52 (s, 1H), 9.24 (d, *J* = 6.1 Hz, 1H), 8.97 – 8.92 (m, 1H), 8.59 (s, 1H), 8.26 (dd, *J* = 8.1, 6.1 Hz, 1H), 8.17 (s, 1H), 7.17 – 7.12 (m, 2H), 6.88 – 6.83 (m, 2H), 4.72 (t, *J* = 7.4 Hz, 2H), 3.73 (s, 3H), 2.63 (t, *J* = 7.8 Hz, 2H), 2.28 (p, *J* = 7.7 Hz, 2H).

$^1$H NMR of 1-(3-(4-methoxyphenyl))propylnicotinamide bromide (P3NA-OMe$^+$ Br$^-$)

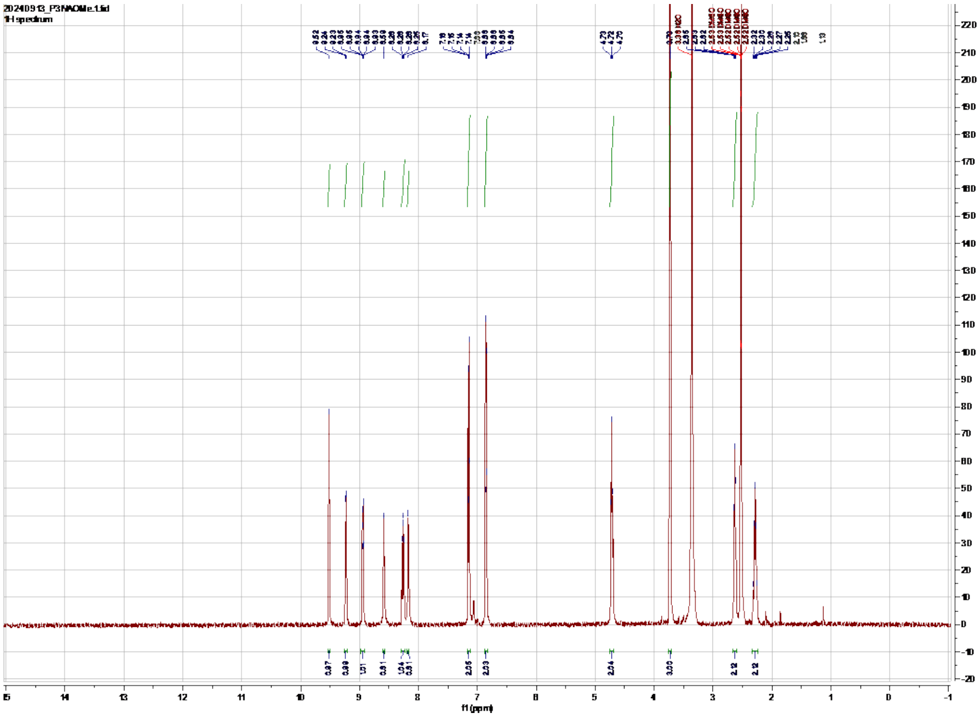


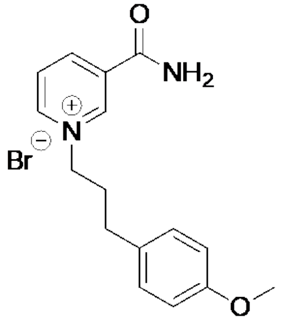

**Synthesis and Characterization of 1-(2-carbamoylmethyl)nicotinamide bromide ($AmNA^+$ $Br^-$)**

1-(2-carbamoylmethyl)nicotinamide (1-(2-amino-2-oxoethyl)-3-carbamoylpyridin-1-ium bromide, $AmNa^+$ $Br^-$) was chemically synthesized using the general procedure reported in King et al[1] with the following starting materials: nicotinamide (750 mg, 5.44 mmole) 2-bromoacetamide (664.33 mg, 5.44 mmole). 1-Bromo-3-phenyl propane was obtained from Sigma. The title compound was isolated as a fine powder in 58% yield (3.157 mmole, 821.5 mg).$^1$H NMR (498 MHz, DMSO) δ 9.46 (d, $J$ = 1.6 Hz, 1H), 9.12 (dt, $J$ = 6.1, 1.3 Hz, 1H), 9.03 (dt, $J$ = 8.2, 1.5 Hz, 1H), 8.61 (s, 1H), 8.32 (dd, $J$ = 8.1, 6.1 Hz, 1H), 8.18 (s, 1H), 8.11 (s, 1H), 7.75 (s, 1H), 5.52 (s, 2H).

1H NMR of 1-(2-amino-2-oxoethyl)-3-carbamoylpyridin-1-ium bromide (AmNa+ Br-)

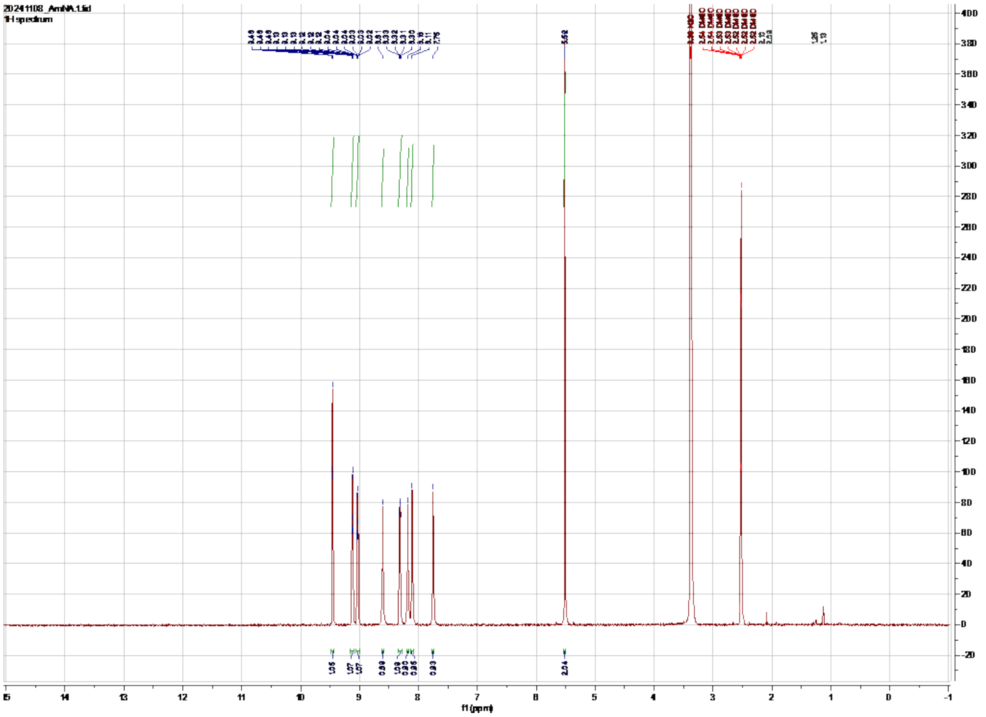


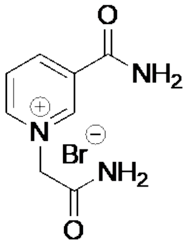


**REFERENCES**


1) Li, H., & Liao, J. C. (2013). Engineering a cyanobacterium as the catalyst for the photosynthetic conversion of CO2 to 1, 2-propanediol. *Microbial cell factories*, *12*(1), 4.
2) King, E., Maxel, S., Zhang, Y., Kenney, K. C., Cui, Y., Luu, E., Siegel, J. B., Weiss, G. A., Luo, R., & Li, H. (2022). Orthogonal glycolytic pathway enables directed evolution of noncanonical cofactor oxidase. *Nature Communications*, *13*(1), 7282.
3) Zhou, J., Gu, X., Zhu, Y., Tao, Z., & Ni, Y. (2023). Engineering glucose dehydrogenase to favor totally synthetic biomimetic cofactors containing carboxyl group. *ChemBioChem*, *24*(15), e202300066.
